# NPAS4 is an allostatic regulator of POMC neuronal activity during diet-induced obesity

**DOI:** 10.1101/2024.01.12.574247

**Authors:** Ji Soo Yoon, Daniel Gamu, William T. Gibson, Francis C. Lynn

**Affiliations:** BC Children’s Hospital Research Institute, Vancouver, BC; Department of Surgery, University of British Columbia, Vancouver, BC; Department of Medical Genetics, University of British Columbia, Vancouver, BC; School of Kinesiology, University of British Columbia, Vancouver, BC; School of Biomedical Engineering, University of British Columbia, Vancouver, BC

## Abstract

**Rationale:** Obesity is characterized by a chronic positive energy balance and altered function of cell types that regulate food intake. These cell types include proopiomelanocortin (POMC) neurons in the arcuate nucleus (ARC) of the hypothalamus that detect peripheral signals and promote a reduction in food intake upon activation. Downstream of neuronal activation, activity-regulated genes such as Neuronal PAS domain protein 4 (Npas4) are induced as part of the response to environmental stimuli. Npas4 is known to have cytoprotective roles in both neurons and pancreatic beta cells. A previous Npas4 knockout study in both mouse pancreatic beta cells and ARC neurons implied a potential role of Npas4 in regulating food intake. However, the specific sites of Npas4 action in the ARC are unknown. We hypothesized that Npas4 in POMC neurons of the ARC has a role in regulating food intake during obesity.

**Methods:** We quantified *Npas4* expression in POMC neurons of the arcuate nucleus in mice exposed to various positive energy states known to activate POMC neurons using RNAscope fluorescent *in situ* hybridization. Next, we generated adult male mice with a conditional Npas4 knockout specifically in their ARC POMC neurons (POMC-NPAS4 KO) and metabolically characterized them for 30 weeks on regular chow or 60% high-fat diet (HFD) at room temperature. In addition, we performed single cell RNA sequencing (scRNA-seq) on microdissected ARC tissue and neighbouring regions from fasted or 1hr refed POMC-NPAS4 KO mice and controls at 6 weeks of HFD, in order to identify Npas4-regulated and feeding-regulated transcriptional changes in POMC neurons.

**Results:** Npas4 was expressed in POMC neurons, and its expression was induced in response to positive energy states such as refeeding, oral glucose, and acute HFD feeding. HFD-fed POMC- NPAS4 KO males showed significantly reduced body weight starting at 10 weeks of HFD, and weighed 8-10 grams less than controls by 30 weeks. With metabolic cages and manual food intake measurements, we determined this difference was not the result of increased energy expenditure or physical activity, but was due to decreased food intake prior to the observed lack of gain in body weight. Using the ARC single-cell dataset, we found that POMC neurons of KO mice showed an enhanced refeeding-induced transcriptional response, dysregulated immediate early gene expression in response to refeeding, and reduced expression of genes encoding GABA-A receptor subunits. Furthermore, cell-to-cell communication analysis revealed that POMC neurons of KO mice specifically lost inhibitory GABAergic signaling inputs, some of which came from agouti-related protein (AgRP) neurons, and gained excitatory glutamatergic signaling inputs compared to POMC neurons of control littermates.

**Conclusions:** Taken together, the results suggest that activity-dependent expression of Npas4 in POMC neurons tempers the activity of these cells upon overnutrition. Loss of Npas4 causes cell-autonomous loss of the capacity to sense nutrient intake. Molecularly, this is driven by reduced expression of inhibitory GABA-A receptors and an overall increase in POMC neuronal activity, leading to decreased food intake and decreased weight gain. In conclusion, for the first time we report a role for the transcription factor Npas4 in POMC neurons of the ARC, and demonstrate it plays an indispensable role in controlling feeding behavior in states of overnutrition.

## Introduction

Mammalian body weight regulation is largely influenced by feeding behaviour, which is controlled by the central nervous system (CNS) [1,2]. In the CNS, there are several distinct regions in the hypothalamus that regulate feeding, including the arcuate nucleus (ARC), ventromedial hypothalamic nucleus (VMH), paraventricular nucleus (PVN), and the dorsomedial hypothalamic nucleus (DMH) [3,4]. In particular, the ARC contains two distinct first-order neuronal populations with opposing functions: an orexigenic population expressing agouti-related peptide (AgRP) [1,5] and an anorexigenic population expressing alpha melanocortin stimulating hormone (α-MSH), which is derived through proteolytic processing of the precursor POMC protein [1,6,7]. Nutrient deficit or hormonal indicators of hunger such as ghrelin activate AgRP neurons to promote food intake [8–10], while indicators of nutrient excess such as elevated glucose, fatty acids, or leptin, activate POMC neurons to promote satiety [11–17].

Ultimately, this system relies on AgRP and POMC neurons secreting the neuropeptides AgRP and α-MSH to inhibit or activate higher-order neurons through the melanocortin-4 receptor (MC4R), respectively [4,18].

In addition to neuropeptide secretion, neuronal activation can be coupled to transcriptional changes through the immediate early genes (IEGs). In response to context-dependent extracellular stimuli, IEGs display a characteristic rapid and transient increase in expression (2- to 10-fold within 1 hour) independent of protein synthesis, and many IEGs encode for transcription factors that regulate downstream target genes for long-term responses [19–21].

Neuronal PAS domain protein 4 (Npas4) is one such IEG that is also a basic helix-loop-helix PAS (bHLH-PAS) family transcription factor, known to be selectively induced in response to neuronal activity and the resulting depolarization with calcium influx [22]. As the name implies, Npas4 was previously shown to be neuron-specific in the CNS across various sites including the cortex, olfactory bulb, and the hippocampus, but also the PVN and ARC of the hypothalamus [22–26]. Npas4 is implicated in a wide range of processes such as neuroprotection, contextual memory formation, and neuronal development [27–31]. At the molecular level, Npas4 activates distinct gene sets in excitatory and inhibitory neurons, regulating inhibitory-excitatory balance within neural circuits, partially through its downstream target Brain-derived neurotrophic factor (Bdnf) [32–34].

Despite the bulk of Npas4 studies being focused on neurons, there have been previous reports of Npas4 in non-neuronal cell types outside of the CNS, most notably the endothelial cells [35] and pancreatic beta cells, where it is also induced in a calcium-dependent manner [36]. One of these extra-neuronal Npas4 studies recently highlighted a potential role in the ARC. In a previous Npas4 knockout mouse line using the tamoxifen-inducible Pdx1-CreER transgene, recombination was observed in both the pancreatic beta cells and regions of the mediobasal hypothalamus (MBH) that included the ARC [37]. When the knockout mice were fed a high-fat diet (HFD), they displayed impaired glucose homeostasis and increased light-cycle food intake [37]. However, a more beta cell-specific Npas4 knockout mouse line using the Ins1-Cre transgene showed no differences in feeding, which implicated Npas4 having a role in ARC neurons that regulate food intake, such as the POMC neurons. To date, Npas4 has not been studied in any ARC neuronal population.

In this study, we specifically examined whether Npas4 has a role in the satiety-promoting ARC POMC neurons in the regulation of food intake. We show that Npas4 is indeed expressed in POMC neurons and can be induced by systemically increasing energy availability to activate these neurons through refeeding after a fast, oral glucose administration, and acute HFD feeding. Surprisingly, we found that conditional knockout of Npas4 specifically in adult POMC neurons in male mice protects them from diet-induced weight gain during chronic HFD feeding. These knockout mice display reduced food intake early in the HFD feeding regime, prior to the significant changes in body weight. To investigate the molecular mechanism behind this phenotype and to identify additional feeding-regulated genes, we conducted scRNA-seq on ARC tissue of fasted and fast-refed control and Npas4 knockout mice. We found that in POMC neurons with Npas4 knockout, there is an enhanced transcriptional response to refeeding but a dysregulation of known IEGs. We also found a reduction in genes encoding for GABA receptor subunits in Npas4 knockout POMC neurons. Further analysis using a cell-cell communication tool showed that there were decreased GABAergic signaling inputs and increased glutamatergic signaling inputs into Npas4 knockout POMC neurons compared to control POMC neurons.

Overall, our findings indicate that under obesogenic conditions, Npas4 acts in allostatic manner to maintain an inhibitory tone on POMC neurons as they are increasingly activated by nutrient stresses, partially through regulating inhibitory GABA receptor expression in these cells.

## Materials and Methods

### Animals

All animal protocols were approved by the University of British Columbia Animal Care Committee. All mice were housed at the Animal Care Facility in the British Columbia Children’s Hospital Research Institute. All experiments and measurements were performed in male mice between 6 to 37 weeks old. Up to four littermates of the same sex were housed per cage with *ad libitum* access to food and water, unless food intake was being monitored. All cages were held in temperature- and humidity-controlled rooms with a 12-hour light/dark cycle at 22°C. All mice were weaned at 3 weeks of age onto standard rodent Chow (Teklad 2918). Mice were switched to HFD (Research Diets D12331) at 7 weeks of age. HFD pellets were refreshed weekly.

Mouse lines used in this study were Npas4^flox/flox^ [22], POMC-CreER transgenics with Rosa26^LSL-tdTomato^ reporter [38], and C57BL/6J strains (Jackson Laboratory strain #000664). All strains were maintained inbred on a C57BL/6J background. For experiments involving the POMC-CreER strain, Npas4^flox/flox^ mice (Npas4 flox) were used as littermate controls, POMC- CreER^+^ (POMC-CreER) mice were used as transgenic controls, and POMC-CreER^+^;Npas4^flox/flox^ mice were used as knockouts (POMC-NPAS4 KO). Ear punch biopsies were taken from mice for PCR genotyping at 3 to 4 weeks of age, after weaning. At 6 weeks of age, 8mg of tamoxifen (Toronto Research Chemicals; T006000) was administered by oral gavage to each mouse every other day, for a total of 3 times over 5 days. Tamoxifen solution was prepared fresh each time by dissolving in corn oil (ACH Food Companies Inc.; 100% pure Mazola corn oil) for a concentration of 60mg/mL.

### Body weight and blood glucose measurements

Body weights of mice were measured weekly using a digital scale between 09:00 and 10:00. Blood glucose was measured biweekly after an overnight fast for 10-12 hours starting at 8 weeks of age. The left saphenous vein was pricked with a 27½ gauge needle (VWR; BD305109) and blood glucose was measured using a handheld OneTouch UltraMini glucometer (LifeScan Europe; AW 06720302A) and OneTouch Ultra glucose strips (LifeScan Europe; AW 06858804A).

### Glucose and insulin tolerance tests

For oral glucose tolerance tests (OGTT), mice were fasted for 10-12 hours overnight before the tolerance test. In the morning, body weight and blood glucose were measured, and one heparinized microcapillary tube (Fisherbrand; 22-362-566) of blood was taken from each mouse (0 minutes). 2g/kg glucose was administered to each mouse by oral gavage, using a sterile-filtered 40% D-glucose (Sigma; G7021-1KG) solution. Blood glucose was measured at 10, 30, 60, 90, and 120 minutes post-gavage, and 1 microcapillary tube of blood was taken at 10 minutes post-gavage.

For insulin tolerance tests, mice were fasted for 2 hours during the light cycle before the tolerance test. Body weight and blood glucose was measured, and 0.75U/kg insulin was administered to mice via intraperitoneal injection with a 27½ gauge needle, using a sterile-filtered 150mU/mL insulin solution in PBS. Blood glucose was measured at 10, 30, 60, 90, and 120 minutes post-injection.

### Plasma collection and ELISA

All blood for plasma collection was collected from the saphenous vein. The blood was transferred to a 1.5mL microcentrifuge tube (Diamed; SPE155-N) and stored on ice for up to 30 minutes before being centrifuged at 5000g for 10 minutes at 4°C. Plasma was collected and transferred to a clean 1.5mL tube. Plasma samples were stored frozen at -20°C until assayed on a rodent insulin ELISA (Alpco; 80-INSMR-CH10) according to manufacturer’s instructions.

### Body composition and metabolic cages

Body composition was measured using an EchoMRI-100H Body Composition Analyzer (Echo Medical Systems). For metabolic caging experiments (LabMaster TSE systems), 13-week-old mice were singly housed and allowed to acclimate within the system for 24 hours before recording for 72 hours. Whole-body energy expenditure was determined via indirect calorimetry by continuously measuring CO2 production rate (VCO2), O2 consumption rate (VO2) and respiratory exchange ratio (RER: VCO2/VO2) throughout the collection period. Food and water intakes was also measured by weight sensors linked to the food hopper and water bottle in each cage, while spontaneous locomotor activity was measured with infrared beam breakage.

### Fast-refeeding and manual food intake measurements

Mice were singly housed and fasted overnight for 10-12 hours by transferring to a clean cage with no food hopper and only *ad libitum* access to water. In the morning, a pre-weighed food hopper with food pellets was placed in the cage for 1 hour. Before and after the refeeding period, body weight was measured for each mouse. If euthanizing for tissue harvest after refeeding, stomach contents were also checked during the dissection to confirm the mouse had eaten.

Weekly food intake was calculated by subtracting the weight of food in the hopper and food crumbs on the cage floor from the weight of the freshly added food given the week before.

### Brain harvest, processing, and sectioning

Whole brains were harvested from euthanized mice following a cold PBS flush and transcardial perfusion with 4% paraformaldehyde (PFA). The brains were post-fixed in 5-10mL of 4% PFA overnight at 4°C, then moved through a sucrose gradient over the next two days before being embedded and frozen in OCT cryomatrix (Epredia; ref 6502). Frozen brains were sectioned in a cryostat (Leica; CM 1950) to obtain 15μm sections at the level of the arcuate nucleus, where the 3^rd^ ventricle was visible. Protocol details can be found in the supplementary methods.

### RNAscope fluorescent *in situ* hybridization

The RNAscope multiplex fluorescent v2 kit (ACDbio) was used according to manufacturer’s instructions for target probes *Npas4* (461631), *Pomc* (314081-C2), and *tdTomato* (317041-C3). A technical positive and negative control was included for every RNAscope experiment using the provided positive and negative control probe mixtures. For development of each target signal, channel-specific HRP reagents and fluorescent Opal dyes 520, 570, or 650 (Perkin Elmer FP1487001KT, FP1488001KT, or FP1496001KT) were used. The mounted slides were imaged with a Leica SP8 confocal microscope and quantified using Imaris v9 (Oxford Instruments).

RNAscope protocol details and quantification can be found in the supplementary methods.

### Immunostaining

Briefly, 15μm fixed frozen sections of mouse brain were washed, permeabilized with 1% SDS, blocked with 5% horse serum in PBS, and incubated overnight with primary antibodies diluted in blocking solution at 4°C. The next morning, the sections were washed with PBS and incubated with secondary antibodies and nuclear stain TOPRO-3 diluted in blocking solution for 1 hour at room temperature. The sections were washed, mounted, and imaged with a Leica SP8 confocal microscope. Protocol details and antibody dilutions can be found in the supplementary methods.

### Mouse hypothalamus dissociation for scRNA-seq

After an overnight fast with or without 1 hour of refeeding in the morning, mice were anesthetized with isoflurane and euthanized by cervical dislocation. The brain was rapidly removed from the skull and the mediobasal hypothalamus area containing the ARC was manually dissected out from each brain. The dissected neural tissue was dissociated according to previously published protocols with some adjustments [39,40], keeping the tissues on ice whenever possible. The dissociated cell solution was filtered, spun down, and resuspended for counting. For the final cell solution, the cells were spun down and resuspended to an appropriate volume of PBS- (cytiva; SH30028.02) for scRNA-seq library preparation. Dissociation protocol details can be found in the supplementary methods.

### scRNA-seq of mouse ARC

Each mouse was processed as an individual sample. Libraries were generated for 10000 target recovery of cells per sample with the Chromium Next GEM Single Cell 3’ version 3.1 reagent kits (10x Genomics) according to the manufacturer’s instructions. Library quality assessment was performed twice during the library generation protocol with the Agilent Bioanalyzer High Sensitivity dsDNA kit (Agilent; 5067-4626). Completed libraries were quantified with both the Qubit high sensitivity dsDNA kit (Thermo Fisher Scientific; Q32854) and KAPA library quantification kit for Illumina platforms (KAPA Biosystems; KK4824). Libraries were sequenced on the Illumina NextSeq500 until all samples reached a minimum of 20000 reads/cell. Sequencing details can be found in the supplementary methods.

### Analysis of scRNA-seq data

Cell Ranger v7.0 (10x Genomics) was used for demultiplexing, aligning to reference genome GRCm38, and counting barcodes and UMI. Outputs of Cell Ranger were carried forward to the R package Seurat [41] v4.0.4 for further filtering, normalization, dimensional reduction, clustering, and downstream analysis/visualization. For identifying DEGs across fasted vs refed states or genotypes, cells expressing *Pomc* were subsetted first, and only cells from the targeted combination of fasted and refed states, POMC-CreER (CT) and POMC-NPAS4 KO (KO), or combinations of the metadata were used in the DEG analysis with *FindMarkers.* Gene ontology (GO) analysis was performed through EnrichR [42–44] and cell-cell communication analysis was performed with NeuronChat [45]. Further details of scRNA-seq analysis can be found in the supplementary methods.

### Statistical Analyses

Statistical calculations were performed in Rstudio or GraphPad Prism v8.0.1. Data are shown as mean ± SEM unless otherwise stated in figure legends. Statistical tests used for each figure are stated in the figure legends. For scRNA-seq data, cutoffs for all statistically significant DEGs were Padjusted < 0.05 using Wilcoxon rank-sum tests. All RNAscope quantifications are shown with each dot representing one biological replicate, as a sum of quantifications from three ARC sections from a biological replicate.

## Results

### Npas4 expression can be induced in POMC neurons in a positive energy state

In order to detect and quantify *Npas4* expression in the ARC POMC neurons, we used the RNAscope FISH technology. We first validated the *Npas4* probe in the hippocampus, a known site of *Npas4* expression [22], and confirmed that the probe is robust and specific (Supplementary Fig.1A). *Npas4* expression is induced in an activity-dependent manner in neurons, and POMC neurons are known to be activated in states of positive energy balance [7,13–16]. Therefore, we hypothesized that acute refeeding or circulating factors that mimic the postprandial state would induce *Npas4* in POMC neurons. Marked differences in *Npas4* expression were observed in ARC sections of adult male B6 mice 1 hour after refeeding following an overnight fast (Fig.1A), in *Pomc+* cells (Fig.1B). In the fasted state, 70% of *Pomc+* cells had detectable *Npas4* expression (1 or more spots/cell), but this proportion was significantly increased after 1 hour of refeeding (Fig.1C). Furthermore, quantifying *Npas4* expression levels as spots per *Pomc+* cell showed significant decreases in the proportions of *Npas4-* and *Npas4*- low *Pomc+* cells (1-3 spots/cell) with corresponding significant increases in the proportions of *Npas4-*medium (4-9 spots/cell) and *Npas4-*high (10 or more spots/cell) populations in the refed state (Fig.1D). In addition, ARC sections from fasted and refed mice were immunostained to determine if refeeding also induced Npas4 protein in POMC neurons. We observed a greater number of tdTomato (tdT)-labelled POMC-CreER lineage-labelled neurons with nuclear Npas4 protein in the 1 hour refed state compared to the overnight fasted state (Supplementary Fig.1B). These results show that 1-hour of refeeding after an overnight fast is sufficient to induce significant *Npas4* in POMC neurons and define Npas4 as an activity marker in this cell type.

**Figure 1:**
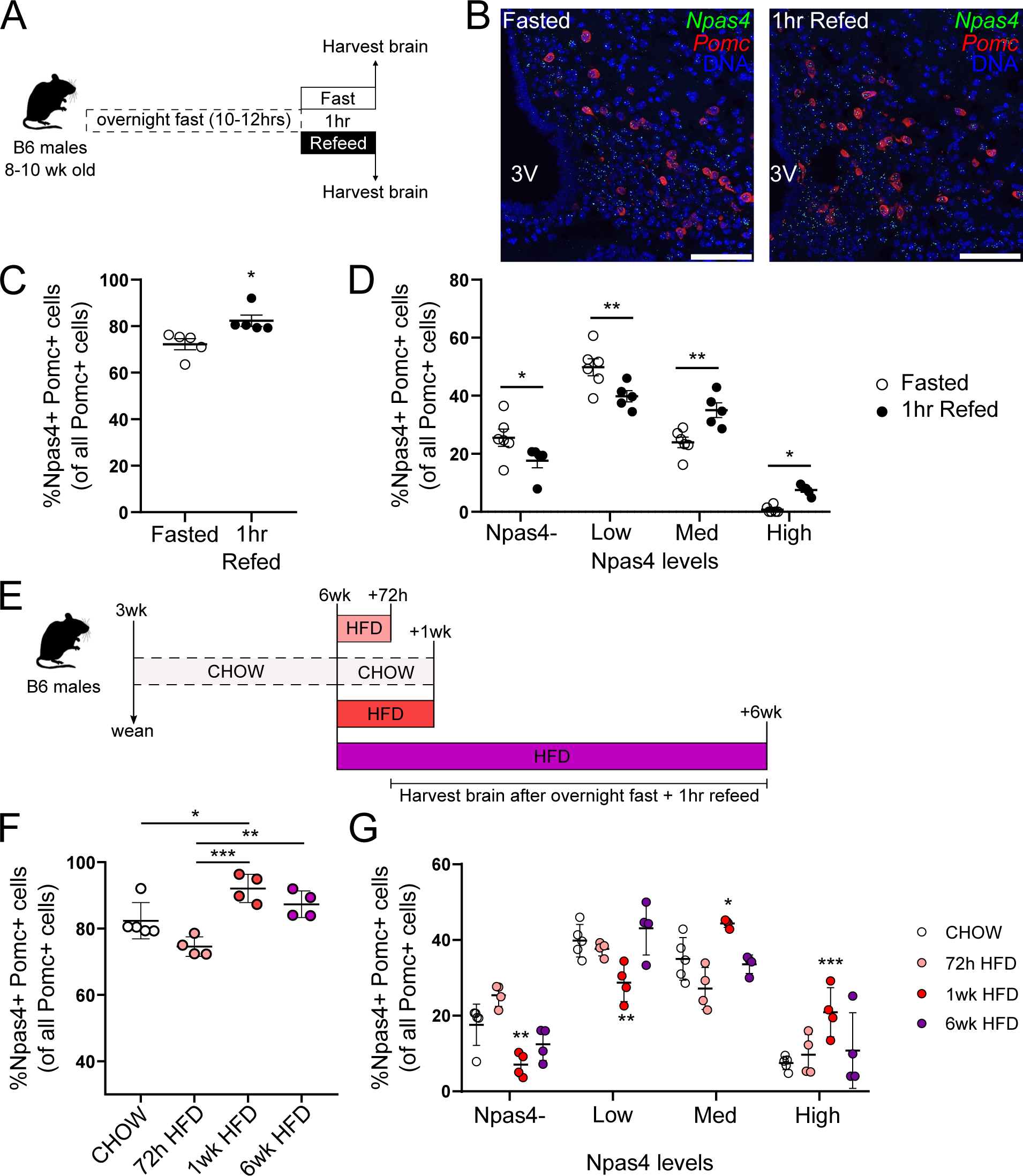
Npas4 is induced in POMC neurons by refeeding. A) Schematic of brain harvests after fasting and refeeding. B) Representative RNAscope images of sections from fasted and 1hr refed mice, probed for *Npas4* (green) and *Pomc* (red) mRNA, with DAPI nuclear stain (DNA; blue). Scale bars = 100μm. C) Quantifications of Npas4+ Pomc+ cells as a percentage of total Pomc+ cells from RNAscope images (B). D) Breakdown of quantifications (C) into bins of Npas4 expression: Npas4- = 0 spots/cell, Low = 1-3 spots/cell, Med = 4-9 spots/cell, High = 10 or more spots/cell. Values are expressed as a percentage of Npas4+ Pomc+ cells in all Pomc+ cells. E) Schematic of HFD brain harvest timeline. F-G) Quantifications (F) and breakdown (G) of %Npas4+ Pomc+ cells in RNAscope images from HFD-fed, 1hr refed mice, in the same manner as C-D. Student’s t-test (C-D): *P < 0.05. One-way ANOVA with Tukey’s multiple comparisons (F-G): *P < 0.05, **P <0.01, ***P < 0.001 vs CHOW.

As some POMC neurons are known to be glucose-activated [13,46], we next assessed whether oral glucose could induce *Npas4.* We applied the RNAscope protocol on ARC sections from mice after an overnight fast and 5, 15, 30, 60, and 180 minutes after oral glucose gavage to detect *Npas4* and *Pomc* expression (Supplementary Fig.2A). *Npas4* induction in *Pomc+* cells was detectable as early as 5-15 minutes post-gavage (Supplementary Fig.2B). Quantification of *Npas4* expression levels per cell showed a significant increase in *Npas4*-high cells and a decrease in *Npas4-*low cells at 15 minutes post-gavage (Supplementary Fig.2C). These results show that oral glucose can rapidly activate POMC neurons and induce *Npas4*, with the peak of induction at 15 minutes, which is within the possible time frame of induction of an IEG like *Npas4* [21].

While an oral glucose gavage will elevate blood glucose levels to beyond that of typical postprandial levels in a mouse, it will also naturally trigger a spike in circulating insulin levels. However, whether POMC neurons are truly activated by insulin is under debate [47–50]. In order to determine whether insulin can induce *Npas4* expression in POMC neurons, we used intraperitoneal insulin injections after a 2 hour fast. Brains from vehicle-injected mice were collected 10 minutes after injection, and from insulin-injected mice 30 and 60 minutes after injection for analyses (Supplementary Fig.2D). Insulin action was confirmed with blood glucose measurements before injections and before euthanasia at each time point (Supplementary Fig.2E). Peripheral insulin administration did not affect the proportion of POMC neurons expressing *Npas4* at any time over 1 hour (Supplementary Fig.2F&G). These results showed that oral glucose, but not the accompanying rise in insulin, induces *Npas4* in activated POMC neurons within 1 hour.

While it is known that POMC neuronal activity is decreased in rodent models of chronic HFD-induced obesity [51], short-term increases in fatty acids have been shown to initially increase POMC excitability [15]. To investigate the effects of acute and chronic HFD feeding on *Npas4* induction in POMC neurons, 6-week-old male B6 mice were fed Chow or HFD for 3 days, 1 week, or 6 weeks (Fig.1E). As above, all mice were fasted overnight and refed for 1 hour before brain collection. Compared to chow-fed mice, 1 and 6-week HFD-fed mice showed a significant increase in the proportion of *Npas4*-expressing POMC neurons (Fig.1F). Notably, *Npas4* expression per *Pomc+* cell was significantly increased only at 1 week following HFD initiation (Fig.1G; Med, High, red dots).

We conclude that Npas4 expression is detectable in ARC POMC neurons, and is induced by refeeding after an overnight fast, oral glucose, and at least 1 week of HFD feeding in male mice. Since the role of POMC neurons is to regulate energy homeostasis following activation by these various stimuli, determining the specific role of Npas4 in these neurons requires a specific model of Npas4 depletion and a robust method of inducing nutrient stress.

### Generation of POMC-NPAS4 KO mice and assessing POMC-CreER recombination

In order to specifically study the role of Npas4 in POMC neurons, we conditionally knocked out Npas4 in 6-week-old male mice with a POMC-CreER transgene [38]. We studied *POMC- CreER^+^; Npas4^wt/wt^*(POMC-CreER), *POMC-CreER^-^; Npas4^flox/flox^* (Npas4 flox) [22], and *POMC-CreER^+^; Npas4^flox/flox^* (POMC-NPAS4 KO) mice. We first validated knockout by directly measuring *Npas4* levels in POMC neurons in POMC-CreER and POMC-NPAS4 KO mice after an overnight fast and 1 hour refeed. In both genotypes, *Npas4* and *Pomc* signals were detectable across the ARC (Fig.2A). However, quantification of images showed a 20-30% decrease in the *Npas4+ Pomc+* cells in the POMC-NPAS4 KO mice (Fig.2B&C). Additional recombination assessment was performed by visualization of the POMC-CreER lineage marker tdT as in Supplementary Fig.1B. Two weeks after tamoxifen administration, tdT^+^ cells (POMC neurons) were visible in the ARC, indicating successful CreER-mediated recombination in the target region (Supplementary Fig.3A). We found no tdT^+^ cells in the nucleus of the solitary tract (NTS), where a smaller population of POMC neurons are known to be present (Supplementary Fig.3B). In addition to assessing tdT protein expression, *tdtomato* mRNA expression in *Pomc+* cells was measured in the ARC, where we detected *tdtomato* expression in 50% of *Pomc+* cells (Fig.2D&E). Altogether, these assessments of POMC-CreER recombination suggest that Npas4 is knocked out of 50% of POMC neurons in the ARC only.

**Figure 2:**
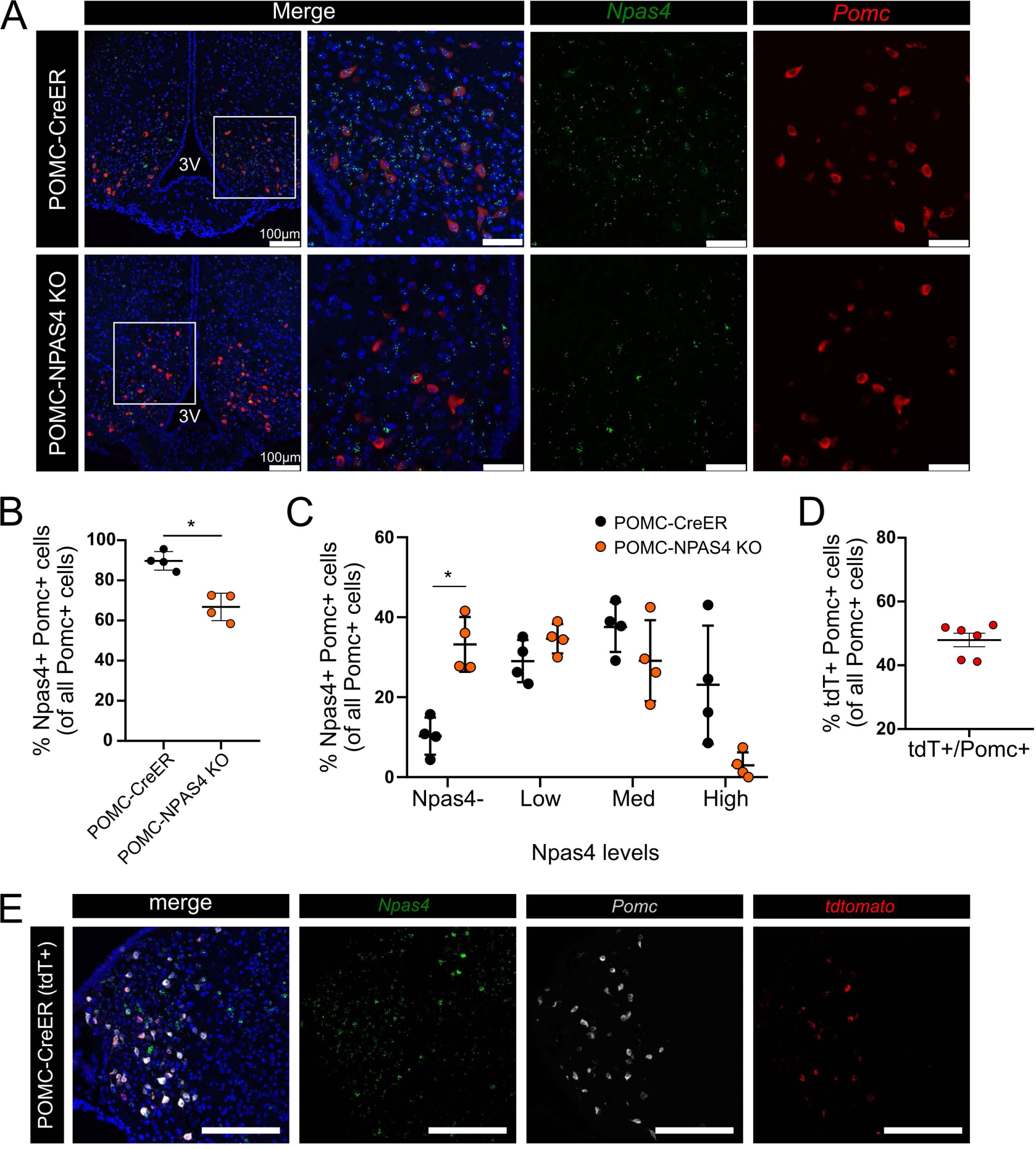
Recombination assessment of POMC-CreER mice. A) Representative RNAscope images of the arcuate nucleus from POMC-CreER and POMC-NPAS4 KO mice, 0 days after tamoxifen administration, probed with *Npas4* (green), *Pomc* (red), and DAPI nuclear stain (blue). B) Quantifications of *Npas4+ Pomc+* cells in each genotype as a percentage of total *Pomc+* cells from RNAscope images (A). C) Breakdown of quantifications (b) into bins of Npas4 expression levels: Npas4- = 0 spots/cell, Low = 1-3 spots/cell, Med = 4-9 spots/cell, High= 10 or more spots/cell. 3V = 3rd ventricle. Data shown as mean± SD. Student’s t-test: *P < 0.05. D) Quantification of *tdtomato+ Pomc+* cells as a percentage of total *Pomc+* cells from E) RNAscope images probed with *Npas4* (green), *Pomc* (white), and *tdtomato* (red) with DAPI nuclear stain (blue). Scale bars= 100µm.

### POMC-NPAS4 KO mice are protected from weight gain during chronic HFD feeding

To characterize the effects of Npas4 KO specifically in POMC neurons, POMC-CreER, Npas4 flox, and POMC-NPAS4 KO male mice were fed either chow or 60% HFD for 30 weeks starting at 7 weeks of age, after tamoxifen administration (Fig.3A). In the chow-fed group, there were no differences in the body weights, fasting blood glucose, glucose tolerance or plasma insulin levels (Supplementary Fig.4A-G). In the HFD-fed group, there were also no differences in fasting blood glucose during 30 weeks of HFD (Fig.3B). However, HFD-fed POMC-NPAS4 KO mice displayed significantly lower body weights compared to both control groups starting at 9 weeks of HFD (Fig.3C). There were no body weight differences before or during tamoxifen administration (Supplementary Fig.5A), and by study endpoint, POMC-NPAS4 KO mice were ∼15% lighter than POMC-CreER and Npas4 flox controls. Glucose tolerance was not different between the three genotypes before (5 weeks), at (9 weeks), and after (29 weeks) the body weight separation occurred (Fig.3D-F). Plasma insulin measured at 0 and 10 minutes during OGTTs was also not different between genotypes throughout the 30 weeks of HFD (Fig.3G-I). Insulin tolerance tests also showed no differences at 10, 18, and 26 weeks of HFD (Supplementary Fig.5B-D). The results of this initial phenotyping of the POMC-NPAS4 KO mice show that the deletion of Npas4 in adult ARC POMC neurons protects male mice from HFD-induced weight gain over time without corresponding changes to their glucose homeostasis.

**Figure 3:**
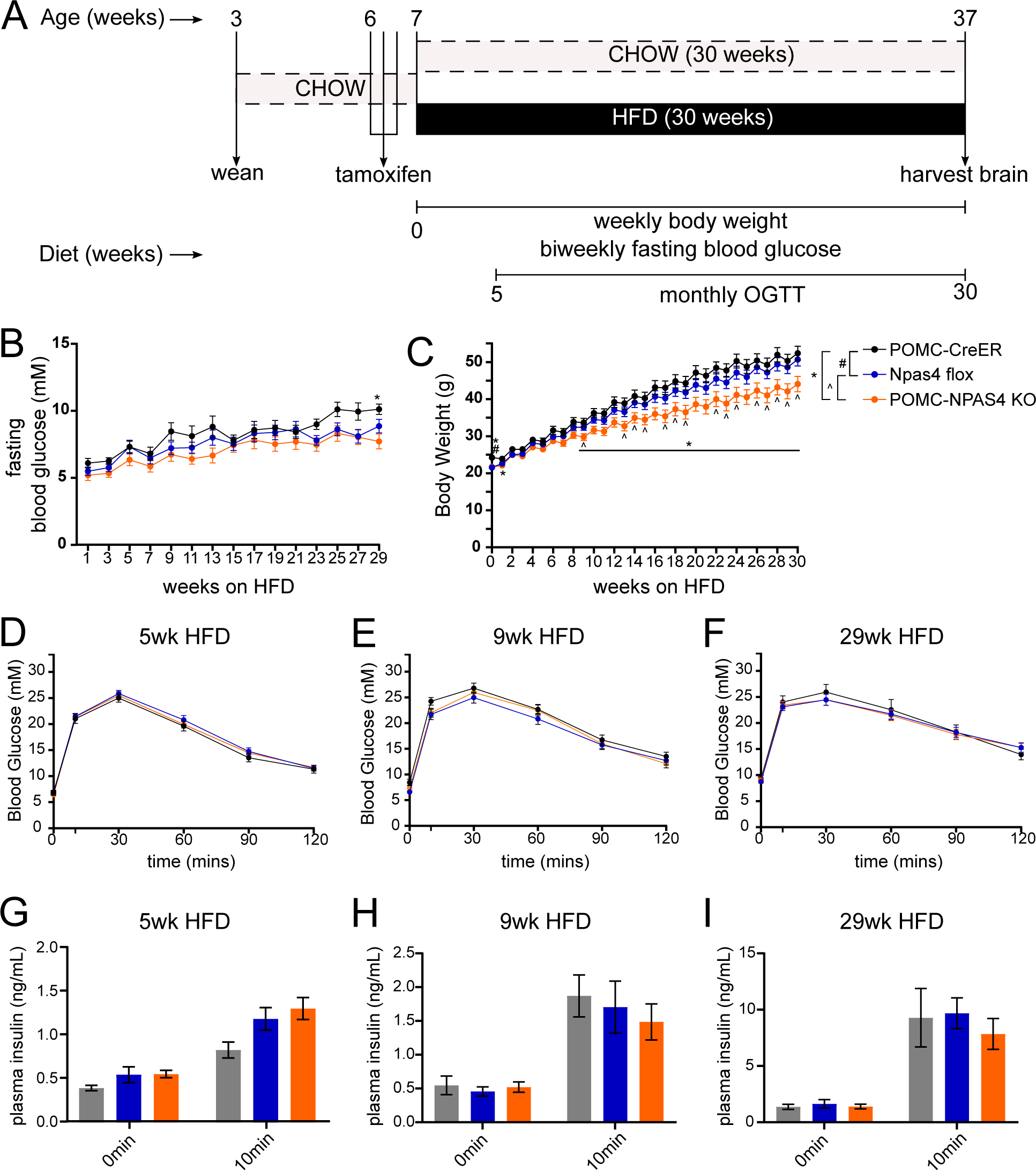
POMC-NPAS4 KO mice gain less body weight on HFD without differences in glycemia. A) Schematic of experiments to characterize POMC-NPAS4 KO mice. B) Biweekly overnight fasted blood glucose measurements for POMC-CreER (n=9), Npas4 flox (n=20), and POMC-NPAS4 KO (n=19) mice during 30 weeks of HFD. C) Body weights measured weekly during 30 weeks of HFD. D-F) Blood glucose measurements during oral glucose tolerance tests at 5 weeks (D), 9 weeks (E), and 29 weeks (F) of HFD. G-I) Plasma insulin measured at 0 and 10 minutes during oral glucose tolerance tests performed at 5 weeks (G), 9 weeks (H), and 29 weeks (I) of HFD. 2-way ANOVA with Tukey’s multiple comparisons: *P < 0.05 (POMC-CreER vs POMC-NPAS4 KO), #P < 0.05 (POMC-CreER vs Npas4 flox), ^P < 0.05 (Npas4 flox vs POMC-NPAS4 KO).

### POMC-NPAS4 KO mice display early reduced food intake during HFD feeding

Due to the known role of ARC POMC neurons in promoting satiety, we hypothesized that POMC-NPAS4 KO mice are protected from severe weight gain on HFD due to reduced food intake. To examine the effects of Npas4 knockout on food intake, activity, and energy expenditure, three POMC-NPAS4 KO mice, three POMC-CreER mice, and two Npas4 flox mice were placed in metabolic cages after 6 weeks of HFD, when the body weight separation had not yet occurred. We observed no differences in oxygen consumption (VO2), carbon dioxide production (VCO2), energy expenditure (estimated as heat production rate) or respiratory exchange ratio (RER), activity or water intake in any genotype (Supplementary Fig.6A-F).

Cumulative food intake over the 72 hours of metabolic caging showed that POMC-NPAS4 KO have reduced food intake compared to Npas4 flox mice (p = 0.0508; Fig.4A). Finally, none of the metabolic cage analyses were affected by differences in fat mass or lean mass, as body composition measurements before metabolic caging showed no differences between genotypes (Fig.4B).

**Figure 4:**
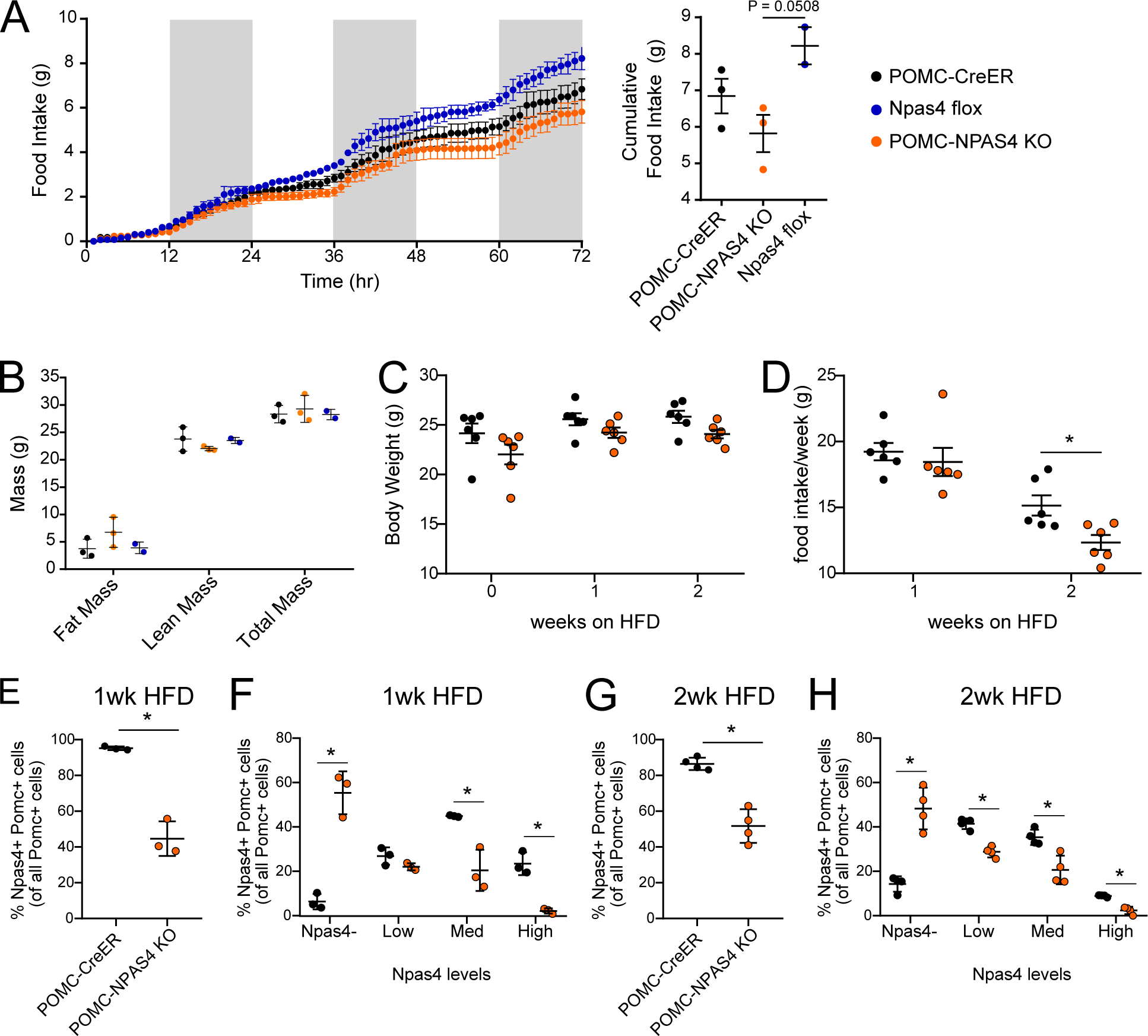
POMC-NPAS4 KO mice show reduced food intake at early weeks of HFD feeding. A) Metabolic cage measurements of food intake (g) for POMC-CreER (n=3), Npas4 flox (n=2) and POMC-NPAS4 KO (n=3) mice over 72 hours, and calculated cumulative total food intake. One-way ANOVA with Tukey’s multiple comparisons. B) Body composition for all mice showing fat mass, lean mass, and total mass before metabolic cages. C) Weekly body weight measurements over 2 weeks of HFD for POMC-CreER (n=6) and POMC-NPAS4 KO (n=6) mice. D) Weekly manual food intake measurements over 2 weeks of HFD. E) Quantifications of *Npas4+ Pomc+* cells from arcuate nucleus sections as a percentage of total *Pomc+* cells from RNAscope images. Brains were harvested after overnight fasting and 1 hour refeeding after 1 week of HFD. F) Breakdown of quantifications (E) into bins of *Npas4* expression levels: Npas4- = 0 spots/cell, Low = 1-3 spots/cell, Med = 4-9 spots/cell, and High = 10 or more spots/cell. G) Quantifications of *Npas4+ Pomc+* cells from arcuate nucleus sections as a percentage of total *Pomc+* cells from RNAscope images. Brains were harvested after overnight fasting and 1 hour refeeding after 2 weeks of HFD. H) Breakdown of quantifications (G) into bins of *Npas4* expression levels as described in F. Data in E-H shown as mean + SD. Student’s t-test: *P <0.05.

We followed up on the results of the metabolic cages by manually monitoring weekly food intake earlier than 6 weeks in the HFD feeding regime. Six POMC-CreER controls and six POMC- NPAS4 KO mice were singly housed for weekly body weight and food intake monitoring.

Consistent with previous data, no body weight differences were observed during the first 2 weeks of HFD feeding (Fig.4C). However, POMC-NPAS4 KO mice showed reduced food intake compared to POMC-CreER mice during the second week of HFD (Fig.4D).

We also assessed Npas4 knockout at both 1 week and 2 weeks of HFD to confirm that the difference in food intake was observed in mice with detectable knockout (Fig.4E). In addition to the increased *Npas4- Pomc+* cells seen at 0 weeks post-tamoxifen previously (Fig.2C), *Npas4-* medium and *Npas4-*high populations were decreased at 1 week of HFD compared to POMC- CreER mice in this cohort (Fig.4F). Similar results were seen at 2 weeks of HFD, although *Npas4-*low *Pomc+* cells were also significantly decreased compared to POMC-CreER mice (Fig.4G&H). These results suggest that the molecular and cellular changes driving reduced food intake happen rapidly upon Npas4 knockout, prior to the reduction in body weights.

### scRNA-seq of fasted and refed ARC of POMC-CreER and POMC-NPAS4 KO mice

In order to identify the molecular and cellular changes in POMC neurons with Npas4 KO we carried out scRNA-seq on the ARC of POMC-CreER (CT) and POMC-NPAS4 KO (KO) mice at 6 weeks of HFD, a timepoint prior to body weight differences that could cause secondary effects. In addition to identifying Npas4-regulated genes, we were interested in identifying other activity-regulated transcriptional changes in POMC neurons. As such, we fasted mice overnight, and refed half of the mice for 1 hour prior to brain collection (Fig.5A). From 7 POMC-CreER mice (4 refed, 3 fasted) and 6 POMC-NPAS4 KO mice (3 refed, 3 fasted), we obtained a scRNA-seq dataset composed of 41,883 cells, integrated between all fasted/fed states and genotypes. This was the sum of 24,399 (58%) KO cells and 17,484 (42%) CT cells, or 16,377 (39%) fasted cells and 25,506 (61%) refed cells. After quality control and filtering steps, the 41,883 cells were clustered and visualized in UMAP space as 33 different clusters (Supplementary Fig.7A). This clustering was minimally impacted by genotypes (Supplementary Fig.7B) but the composition of clusters 7, 14, 22, and 29 showed skewing towards either fasted or refed cells (Supplementary Fig.7C).

**Figure 5:**
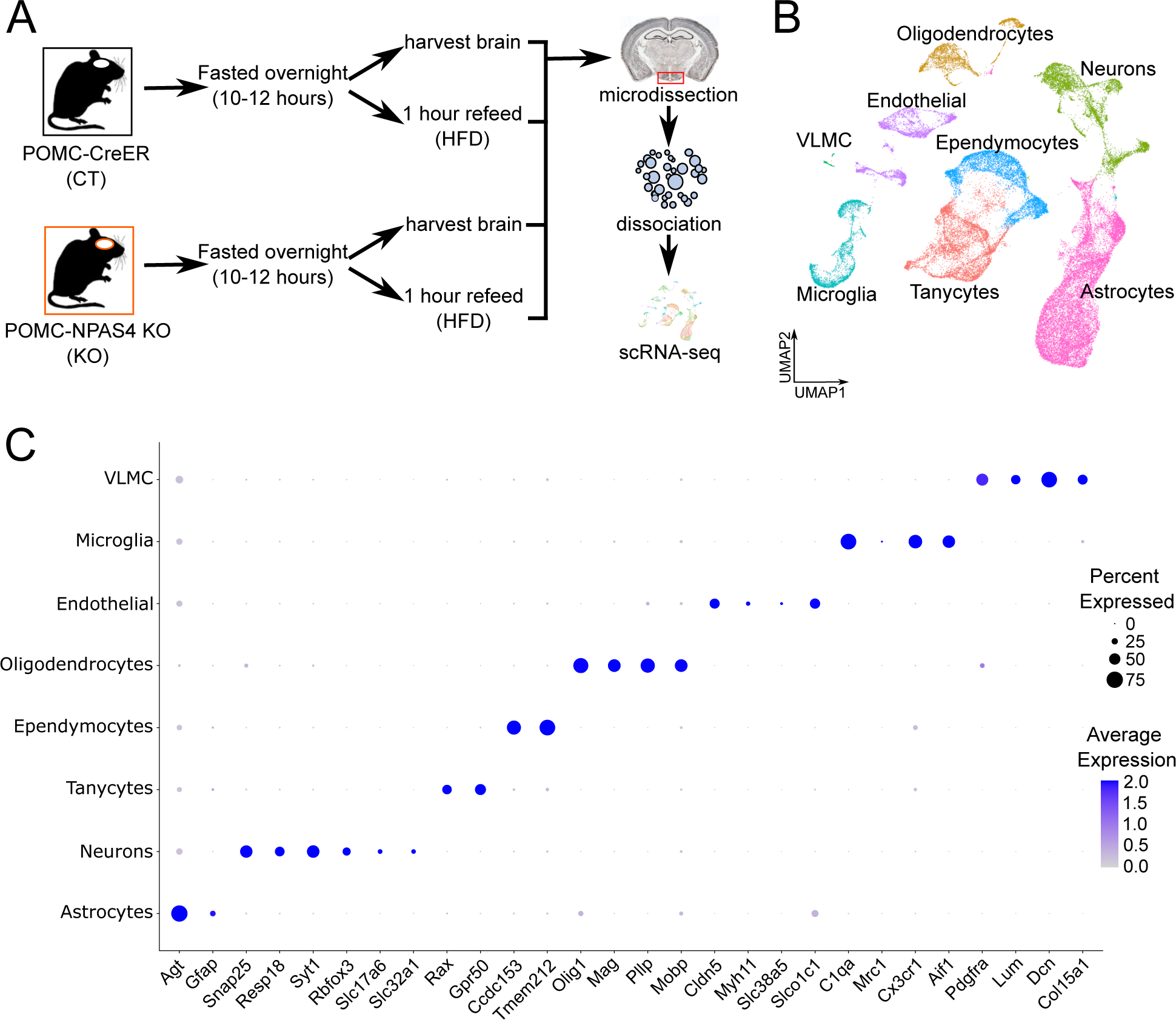
scRNA-seq of mouse ARC cells to identify feeding-regulated genes and NPAS4- regulated genes at 6 weeks of HFD. A) Overview of experimental design to generate single-cell libraries from POMC-CreER (CT; n=3 fasted, n=4 refed) and POMC-NPAS4 KO (KO; n=3 fasted, n=3 refed) mice at 6 weeks of HFD. B) .All cells integrated from all samples and projected in UMAP space, labelled by biological cell types. C) Dot plot showing the average expression in each cluster for the top differentially expressed genes (DEGs) expressed in each cell type. Dot sizes represent percent of cells expressing a gene on the x-axis (“Percent Expressed”) and shade of blue represents expression levels in the cell type (“Average Expression”), with darker blue being higher average expression.

Next, we used known marker genes to biologically classify each of the 32 clusters into a cell type: neurons (*Syt1, Snap25*), oligodendrocytes (*Olig1*), astrocytes (*Agt*, *Gfap*), ependymocytes (*Ccdc153, Tmem212*), tanycytes (*Rax, Gpr50*), endothelial cells (*Cldn5, Myh11*), microglia (*C1qa, Cx3cr1*), and vascular leptomeningeal cells (VLMC; *Pdgfra, Lum*). It became apparent after examining marker genes that the separation of clusters was largely based on cell type (Fig.5B). We also performed an unbiased DEG analysis for each cell type to check if the top DEGs in each cell type overlapped with known marker genes (Supplementary Table 1). For each cell type, we found that 2-4 DEGs were sufficient to confirm the biological identities (Fig.5C).

### Identifying transcriptional impacts of Npas4 KO in POMC neurons after 6 weeks of HFD

A benefit to scRNA-seq is that it allowed us to isolate the relatively rare population of POMC neurons in the computational space. Of the 5,669 neurons, 1,452 *Pomc-*expressing cells were subsetted (Fig.6A). Initially, a general comparison of KO and CT POMC neurons was performed to identify any major transcriptomic changes (Supplementary Table 2). 605 genes were downregulated in KO cells, but only four genes were upregulated in KO cells: *Cwc22, Mt2, Agt,* and *Slc6a11.* All 605 genes downregulated in KO POMC cells were used for a gene ontology (GO) analysis. The majority of the top 20 results from the Molecular Function 2021 library were terms related to neurotransmitter signalling and ion channel activity, including: “transmitter-gated ion channel activity”, “neurotransmitter receptor activity involved in postsynaptic membrane potential”, and “ligand-gated anion channel activity” (Fig.6B). Furthermore, there were several terms that specifically related to GABA signalling: “GABA-gated chloride ion channel activity”, “GABA-A receptor activity”, and “GABA receptor activity” (Fig.6B). In line with the GO results, genes encoding for GABA type A receptor subunits were part of the downregulated gene set in KO POMC cells: *Gabrb3, Gabrb1, Gabrg3, Gabra2, Gabrg1, Gabrg2, and Gabra3* (Fig.6C). In line with this, we observed a robust reduction in GABA type A receptor subunit β3 (GABAAR β3, encoded by *Gabrb3*) protein across the entire ARC region of POMC-NPAS4 KO mice compared to that of POMC-CreER mice at 6 weeks of HFD (Fig.6D).

**Figure 6:**
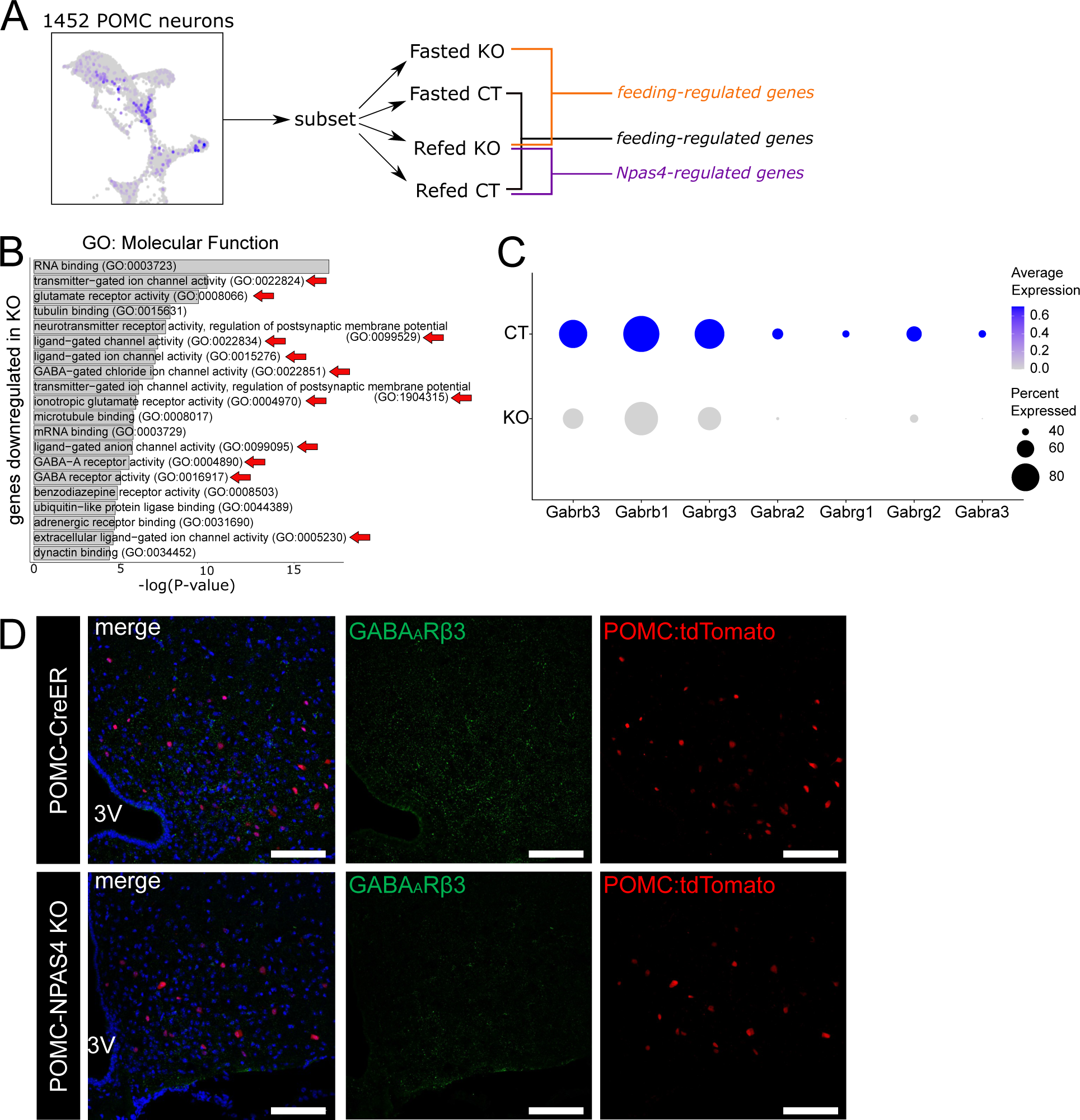
POMC neurons of 6 week HFD-fed POMC-NPAS4 KO mice have reduced GABA receptor subunit expression. A) Schematic showing the analysis strategy used to identify DEGs between genotypes and conditions specifically in subsetted 1452 POMC neurons. DEG analysis was performed for pairs of different genotype and condition combinations to identify feeding-regulated genes and NPAS4-regulated genes. B) Bar plots of the top 20 GO terms from GO library Molecular Function 2021 that are associated with genes downregulated in KO POMC neurons, in order of significance. Red arrows indicate terms associated with neurotransmitter signalling and neurotransmitter receptors. C) Dot plot of significant DEGs enriched in CT vs KO *Pomc+* neurons encoding for GABA-A receptor subunits. The size of dots represents the percent of cells in the genotype that express the gene (“Percent Expressed”) and the shade of blue represent the average expression of the gene on a normalized relative scale, with darker shades of blue representing higher expression levels. D) Representative images of 6 week HFD-fed POMC-CreER and POMC-NPAS4 KO mice, with tdTomato (red) marking POMC neurons in the arcuate nucleus and immunostained for GABA-A receptor beta 3 subunit (green). Scale bars = 100um. 3V = 3rd ventricle.

Taken together, the comparison of KO and CT POMC neuron transcriptomes, regardless of feeding state, showed that the KO neurons displayed reductions in GABAAR genes, and GO analysis suggested there were reduced neurotransmitter and ion channel activities. A reduction in the ability of the Npas4 KO POMC neurons to receive the inhibitory GABA signals could lead to elevated function or activity, which would explain the reduced food intake phenotype.

### Npas4 KO in POMC neurons leads to dysregulation of feeding-regulated transcriptional response

Next, the fasted and refed POMC neurons were compared within each genotype to identify genes that were induced by refeeding. DEG analysis was performed across conditions in CT and KO POMC neurons separately, to generate lists of upregulated genes in the fasted and refed conditions (Supplementary Table 3). In CT POMC neurons, only seven genes were feeding-induced and 88 genes were expressed higher in the fasted condition. KO POMC neurons, on the other hand, showed a much greater transcriptional response in both conditions with 84 feeding-induced and 357 feeding-inhibited genes (Supplementary Fig.8A). All seven of the feeding-induced genes in the CT (*Cdk8, Camk1d, Gm20594, Lars2, mt-Nd3, Luzp2,* and *Prex2*) were also feeding-induced in the KO (Fig.7A). Since we established that refeeding induces *Npas4* in POMC neurons, and refeeding leads to hundreds of transcriptional changes in the KO but not in the CT (Supplementary Fig.8A), the differences in gene expression in the refed condition was expected to reveal Npas4-regulated transcriptional changes. The pairwise comparison revealed 530 genes that were downregulated in KO cells, and only 19 genes that were upregulated in KO cells upon feeding. As expected, the majority (16 out of 19) of these upregulated genes were also detected as refeeding-induced previously. Notably, several well-known IEGs were found to be refeeding-inhibited in KO cells, including *Junb*, *Egr1,* and *Fosb* (Fig.8B). Since IEGs are expected to be upregulated when cells are activated in response to stimuli and the number of activity-regulated genes correlates with cell function in other cell types [52], these findings suggest that KO POMC cells may be more susceptible to greater activity-regulated transcriptional changes, but this response may be dysregulated compared to CT POMC neurons.

**Figure 7:**
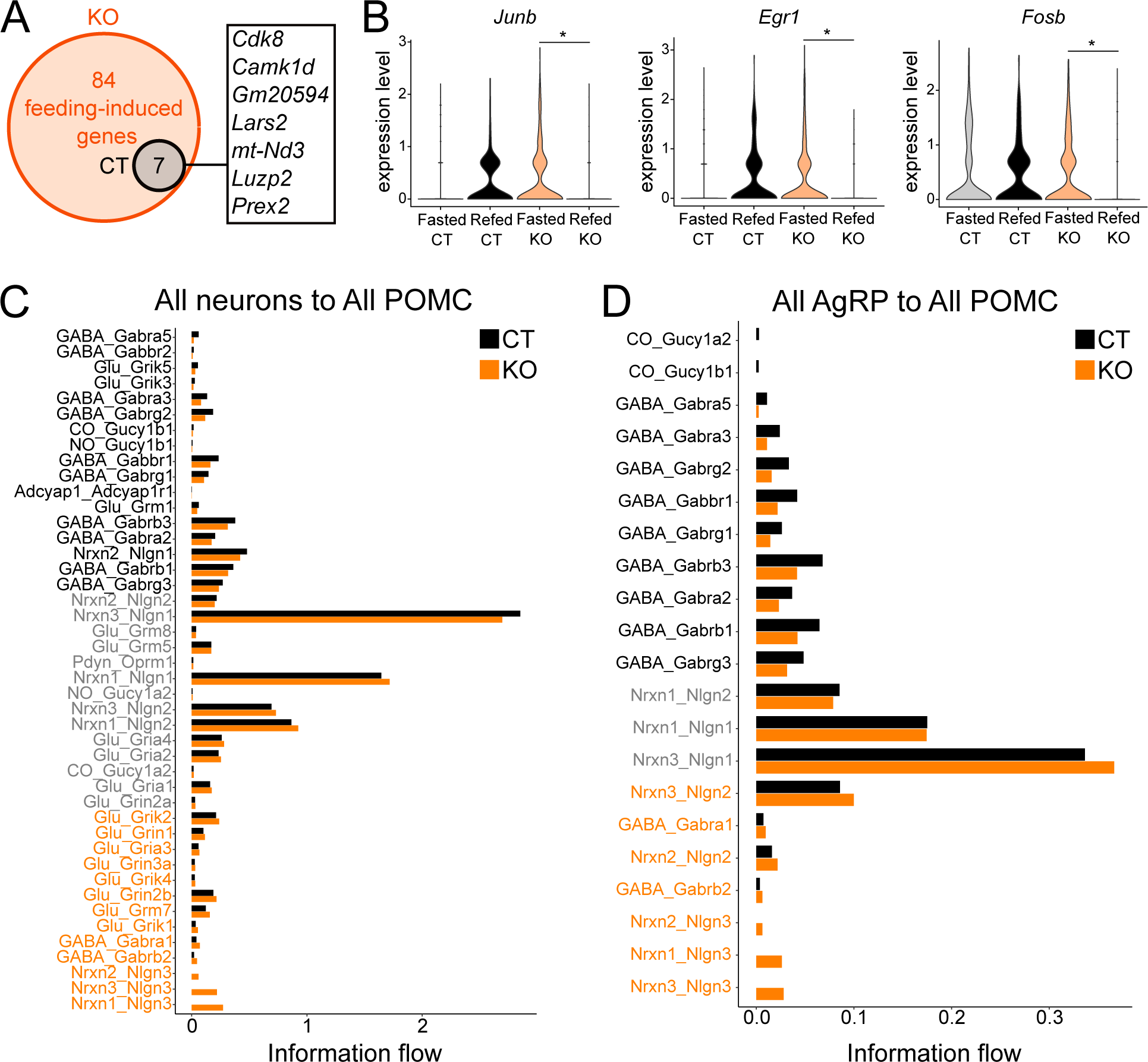
POMC neurons of POMC-NPAS4 KO mice have dysregulated feeding-regulated gene expression and altered GABAergic and glutamatergic inputs. A) Venn diagram showing all seven genes that are induced by refeeding in the CT POMC neurons are also induced by feeding in the KO POMC neurons. The identities of the seven genes are shown in the black box to the right. B) Violin plots showing expression of known immediate early genes that have significantly reduced expression in response to refeeding in POMC neurons of POMC-NPAS4 KO mice but not POMC-CreER controls. Wilcoxon rank-sum test: *P_adjusted_ < 0.05. C-D) Barplots comparing the weight of individual signaling interactions between CT and KO, restricted to signals coming from all neurons to POMC neurons (C) or AgRP neurons to POMC neurons (D). Significantly upregulated interactions in CT cells are labelled in black and significantly upregulated interactions in KO cells are labelled in orange. Grey labels indicate interactions that were not significantly changed between CT and KO.

### POMC neurons with Npas4 KO are predicted to lose GABAergic inputs and gain glutamatergic inputs

Due to the observed downregulation of inhibitory GABA receptors on POMC neurons of KO mice, we decided to computationally investigate if there were additional changes to other neurotransmitter receptors and signaling pathways using a recently published neuron-specific cell-to-cell communication analysis package, NeuronChat [45]. To do this, we first subsetted the 5,669 neurons in our scRNAseq dataset and re-clustered them into 21 clusters (Supplementary Fig.8B). Next, we classified clusters as glutamatergic (*Slc17a6*), GABAergic (*Slc32a1, Gad1*), or mixed (Supplementary Fig.8C). We identified 6 clusters as glutamatergic, 8 clusters as GABAergic, and 7 clusters as mixed. Marker genes that were known to be expressed in neuronal populations within the ARC and other nearby hypothalamic nuclei were used to classify neuron clusters. For ARC neurons, we used the pan-ARC marker gene *Tbx3* [53], as well as *Pomc, Cartpt, Agrp, Npy, Tac2,* and *Pnoc.* We found that there were 2 clusters of both POMC/CART and AgRP/NPY neurons, but AgRP neurons clustered very clearly (Supplementary Fig.8B, 8D). POMC neurons, on the other hand, were more intermingled near the *Tac2*+ and *Pnoc+* neurons. For regions and neuronal populations outside of the ARC, we identified 2 clusters of *Adcyap1+/Thy1+* (PACAP) neurons, a cluster of *Nr5a1+/Fezf1+* (SF1) neurons from the VMH, a cluster of *Pdyn+/Ppp1r17* neurons from the DMH [54,55], and 3 clusters of *Rgs16+* neurons from the SCN that had differential expression of *Vip, Cck,* and *Avp* (Supplementary Fig.8D).

Interestingly, cluster 21 highly expressed *Flt1*, an endothelial marker. In addition, most of the mixed clusters and cluster 2 lacked a significant number of DEGs that could be used to identify their biological nature, which was most likely due to the fact that they were clusters that had lower number of genes overall. These 7 clusters (0, 2, 3, 5, 6, 11, and 12) were left included in the dataset for ongoing analysis while labelled as “undefined”.

We used NeuronChat to identify which neural signaling inputs/outputs were changed between KO and CT cells, with a particular focus on the POMC neurons as the targets of signaling coming from other neuronal populations. Similar to scRNAseq DEG analysis at the level of all POMC neurons, the NeuronChat results showed that KO POMC neurons had significantly downregulated GABAergic signaling inputs that implicated 10 different GABA receptor subunit genes, as well as PACAP signaling inputs (Adcyap1_Adcyap1r1), CO, and NO signaling inputs (Fig.7C). The GABA_Gabra5 signaling interaction showed a particularly robust downregulation in the KO compared to the CT, with inputs from four different clusters predicted to have been lost on one of the POMC neuron clusters specifically (Supplementary Fig.9A). In contrast, there was significant upregulation of neurexins 1-3 signaling through neuroligin 3 and glutamatergic signaling inputs through 8 different receptor genes (Fig.7C). For example, the glutamatergic signaling interaction Glu_Grin2b from the PVN SIM1 neurons to one of the POMC neuron clusters was increased in the KO compared to the CT (Supplementary Fig.9B). Since it is known that AgRP neurons are GABAergic and directly inhibit POMC neurons [56], we also identified significantly different signaling pairs specifically for AgRP neurons set as the “senders” and POMC neurons set as the “targets”. We found that once again, the majority of the downregulated inputs onto KO POMC neurons from AgRP neurons were GABAergic, and the GABA receptor genes included all 7 of the significantly downregulated GABAAR subunit genes in Fig. 6C (Fig.7D). There were also upregulated neurexin-neuroligin signaling in KO POMC neurons, including neurexins 1-3 and neuroligins 2-3 (Fig.7D).

In summary, our scRNA-seq data show that POMC neurons with Npas4 KO are more excitable, likely driven by decreased GABAergic receptor presence and increased glutamatergic receptors and are displaying dysregulated hyperactivity leading to reduced food intake.

## Discussion

In cell types like ARC POMC neurons that must constantly detect and respond to environmental changes, IEGs are essential for coupling the stimuli to long-term changes. In both the healthy state and metabolically diseased states such as obesity, there is value in understanding how the transcriptional response of these cells help them functionally adapt to ongoing challenges. In this study, we demonstrated that Npas4 is expressed in POMC neurons of adult male mice and can be induced by canonical activators of POMC neurons, including refeeding after a fast, elevated blood glucose, and HFD feeding. We showed that adult male mice with Npas4 knockout specifically in POMC neurons gain less body weight over time during chronic HFD feeding due to an early reduction in food intake. We used scRNA-seq on fasted and refed mouse ARC cells and showed that POMC neurons with Npas4 knockout have a dysregulated but hyperactive transcriptional response to refeeding, reduced GABA receptor expression and signaling inputs, and increased glutamatergic signaling inputs. Overall, we have shown that Npas4 acts in an allostatic manner to regulate POMC neuron function and activity-regulated transcriptional response during the development of diet-induced obesity.

It is known that in chronic HFD models of rodent obesity, the activity of POMC neurons is drastically altered and they display decreased basal excitability and firing rate [51,57]. However, acute HFD has been shown to increase *Pomc* mRNA expression and cause synaptic changes in the ARC that results in increased POMC neuronal activity [58–60]. Therefore, we can speculate that during the early days of HFD exposure, POMC neurons are initially hyperactivated to compensate for the sudden increase in nutrient stresses, but ongoing exposure and stimulation eventually exhausts the neurons and renders them dysfunctional during overt obesity. In line with this, we did observe the highest refeeding-induced *Npas4* expression levels in POMC neurons at 1-2 weeks of HFD. After 1 week of HFD feeding, there was not only a significant increase in the proportion of *Npas4^+^* POMC neurons after refeeding, but there was also a notable increase in POMC neurons that expressed medium-to-high levels of *Npas4* (Fig.1F&G). Interestingly, after 6 weeks of HFD, there was a reduction in this population of POMC neurons that expressed higher levels of *Npas4.* Furthermore, during the earliest weeks of HFD feeding, there is a 10% increase in *Npas4-* and *Npas4-*low populations and a corresponding 10% decrease in the *Npas4-* medium and *Npas4-*higher populations of POMC neurons at 2 weeks of HFD compared to 1 week of HFD (Fig.4F, H). From these results, we can conclude that during the course of diet-induced obesity in a normal mouse, Npas4 expression is initially also upregulated as POMC neurons become hyperactive to meet the demands of switch in diet but is reduced over time as the HFD is maintained. Whether the reduction in Npas4 levels over time is a contributing factor to the reduced activity of POMC neurons or a consequence is unclear, but we can appreciate that ARC Npas4 levels are lower during obesity than during leanness.

Notably, the POMC-NPAS4 KO mice reported herein are a genetic model of reduced Npas4 levels prior to HFD diet, and their phenotype of reduced food intake and body weight suggest the opposite of the natural Npas4 levels during the course of diet-induced obesity. After all, if lower Npas4 levels are associated with an obese state, then our POMC-NPAS4 KO mice with reduced Npas4 prior to HFD should have developed obesity more quickly than control mice. In our model, the Npas4 knockout effectively sets the reduced Npas4 levels as a cause rather than a consequence of altered POMC neuronal activity, and shows that unlike normal progression of obesity, the reduced Npas4 levels from the beginning of HFD is actually beneficial due to the early hyperactivity of POMC neurons. In other words, the genetic knockout of Npas4 specifically in POMC neurons does not mimic the normal progression of diet-induced obesity, where the reduction in Npas4 is most likely due to the reduced activity of POMC neurons over time.

To add another layer of complexity, we also observed Npas4 expression throughout the ARC cells, not just POMC neurons (Fig.1B). It is possible that Npas4 carries out different roles and targets different genes in AgRP neurons or other neuronal populations and even subpopulations within the ARC, as it has been shown before that the targets of Npas4 can be different according to the neuron type [32]. Depending on what the outcomes of Npas4 action in other neuronal populations are, we can speculate that some of these outcomes may contribute to the reduced activity of POMC neurons and in turn, reduced Npas4 levels during obesity. Again, this is a very different scenario than our genetic knockout model, where Npas4 is artificially deleted in only a percentage of POMC neurons.

A caveat to almost all Npas4 knockout experiments performed in this study is the variability and efficiency of the POMC-CreER-mediated recombination. The highest recombination rate observed was around 50% at 1-2 weeks of HFD feeding (Fig.4E, G). This was supported by the *tdtomato* detection in 50% of POMC neurons (Fig.2D&E). Even though the POMC-NPAS4 KO mice showed reduced body weight gain on HFD over time, it was a moderate difference compared to the obese controls and occurred weeks into the HFD regime (Fig.3C). Part of this could have been due to incomplete recombination, and remaining POMC neurons with intact Npas4 could have compensated for the neurons with Npas4 knockout, causing a delay in the appearance of phenotype. It is possible that if recombination rates were higher, the mice would have exhibited even less weight gain on HFD. Furthermore, we do not whether the existence of different POMC subpopulations may have impacted Npas4 knockout, and how this impacted the phenotype. POMC neuron heterogeneity is well-documented, with different subpopulations suspected to respond to different signals to contribute to multiple metabolic functions [61].

Naturally, we suspect that there may be significant heterogeneity in both CreER and Npas4 expression across the POMC population as well. Across the various stimuli to induce Npas4, we consistently observed that 10-20% of the detected POMC neurons do not express any levels of *Npas4* within 1 hour (Fig.1D). This could represent a population that is either particularly rapid at downregulating the induced *Npas4* transcript within the hour, or a population that is not responsive to the stimuli we tested. If there were methods to specifically target a more easily excitable subpopulation of POMC neurons for Npas4 knockout, we may have observed a stronger phenotype.

Using our scRNA-seq dataset, we determined that POMC neurons with Npas4 knockout are likely in a dysfunctional but hyperactivated state. The data suggest that Npas4 is normally important for decreasing POMC tone, and the knockout of Npas4 results in increased activity of POMC neurons due to reduced inhibitory and increased excitatory inputs. In the hippocampus, Npas4 is already well-characterized to be important in maintaining inhibitory tone in neuronal circuits by regulating the expression of GABA receptor genes [22]. In the ARC, AgRP neurons are known to directly inhibit POMC neurons by GABAergic signalling [56], and our analysis showed a decreased GABAergic signaling from AgRP to POMC neurons (Fig.7D). Why does an activity-regulated mechanism to inhibit the activity of satiety-promoting neurons exist? From an organismal perspective, rewiring neural circuits and having frequent hyperactivation of POMC neurons, even during conditions of plenty, cannot be favourable in the long term. There is a possibility of exhausting the neurons and making them more susceptible to stress and death over time. Also, the possibility of constantly feeling sated and being prevented from eating clearly would not have been beneficial along the evolutionary timeline, particularly when obtaining food would have been more of a challenge than it is today. However, there is clearly an early compensatory mechanism that rewires the ARC to increase the anorexigenic tone in a HFD setting. During this early time point when POMC neurons are stimulated and adapting to the increase in nutrient availability, Npas4 could be a key factor that acts in an allostatic manner, predicting the ongoing nutrient stress and reducing neuronal activation and eventual exhaustion of the cells by increasing inhibitory receptors. In this way, Npas4 could be important both as a neuroprotective factor but also through regulating synaptic inhibitory tone, as has been studied in other cell types [29,30,32,36]. In addition, since AgRP neurons have also been suggested to fit the model of allostasis rather than homeostasis [62], it seems reasonable to expect that POMC neurons would also fit this model, and Npas4 and other IEGs could be essential in coupling the environmental changes to long-term predictive responses to perceived future demand.

In summary, here we demonstrate for the first time that the IEG transcription factor Npas4 is expressed in ARC POMC neurons and has a role in regulating food intake by regulating genes that can impact POMC neuronal function and activity. These findings have shown the importance of studying activity-regulated genes in energy homeostasis and energy allostasis to further our understanding of how ARC neurons respond in a cell autonomous manner to intermittent and ongoing stimuli in an obesogenic environment.

## Author Contributions

JSY and FL designed the overall research. JSY and DG performed experiments and analyzed data. JSY and FL wrote the manuscript. All authors subsequently read and edited the manuscript. WTG provided essential materials and infrastructure.

## Acknowledgements

We sincerely thank the members of the Lynn lab for their helpful discussions and technical support. We thank Michael Greenberg for providing the Npas4 floxed mouse line and Joel Elmquist and Chen Liu for providing the POMC-CreER line and Syann Lee for technical advice. We thank the late Denis Richard for sharing the POMC-CreER mouse line. We thank Jacqueline Quandt and Pierre Becquart for the kind gift of the NPAS4^-/-^ mouse brains used for RNAscope probe validation.

We are grateful to the Canadian Institutes of Health Research (MOP 142222) for providing operating support for this work. Research infrastructure was funded by the Canada Foundation for Innovation (#33644). Salary support (FCL) was from the BCCHRI, Diabetes Canada, and the MSFHR (BIOM 5238). Research salary support to WTG was provided by the BC Children’s Hospital through an intramural IGAP award. Fellowship support to DG was provided by the MSFHR (#18533). The funders had no role in study design, data collection and analysis, decision to publish, or preparation of the manuscript.

## Competing Interests

The authors declare no competing interests.

---

**Supplementary Figure 1:**
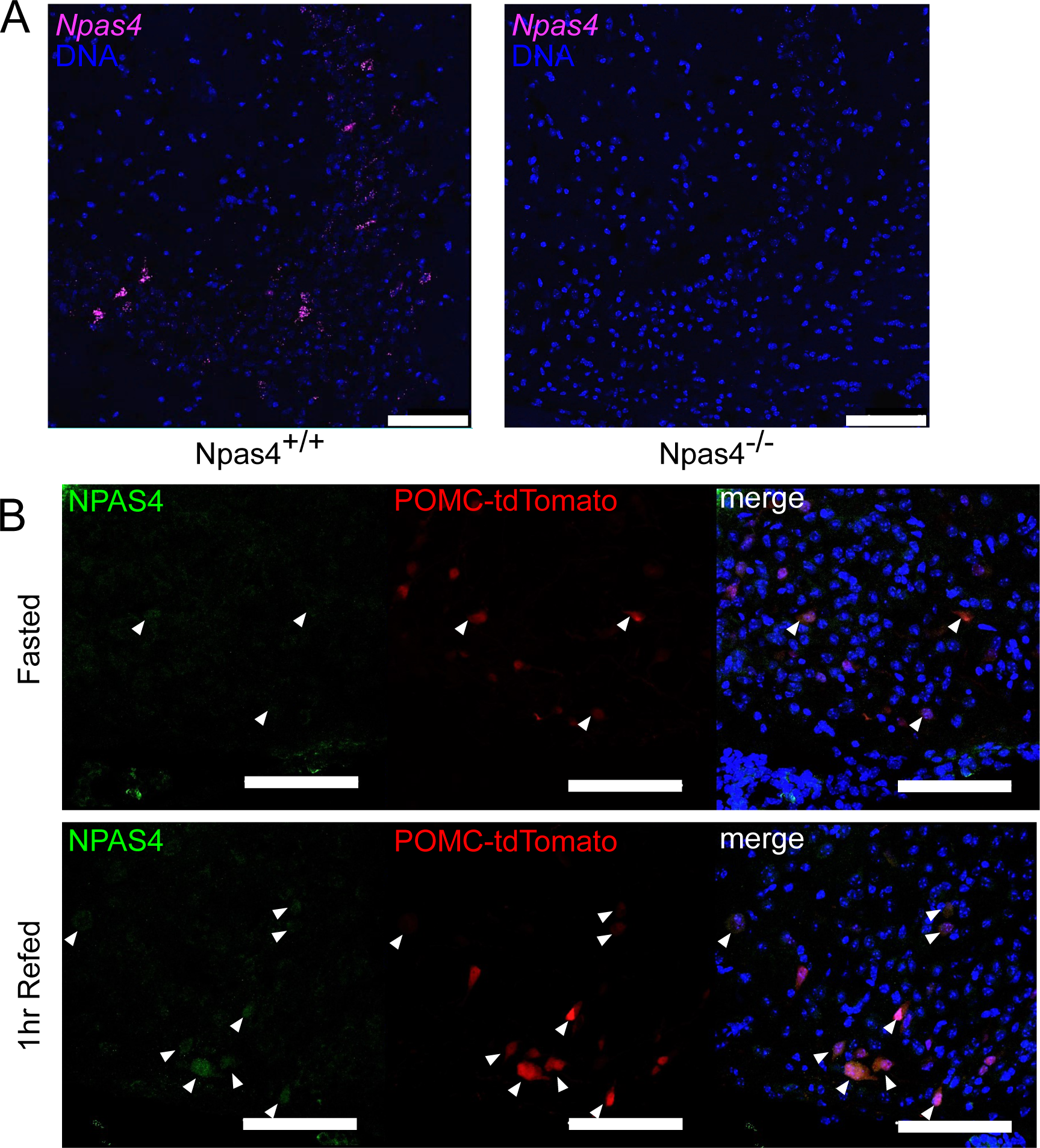
**Detection of Npas4 mRNA and protein**. A) Validation of the *Npas4* RNAscope probe in representative sections of the hippocampus CA3 region from Npas4 +/+ and Npas4 -/- mice. Magenta = *Npas4*, blue = DNA. B) Representative NPAS4 immunostaining images in arcuate nucleus sections of overnight fasted POMC-CreER mice, with or without 1hr refeeding. Brains were harvested 0 weeks after tamoxifen administration to induce expression of nuclear tdTomato protein specificially in POMC neurons. White arrowheads mark cells with co-localized NPAS4 and tdTomato. Green = NPAS4, red = tdTomato labelling POMC neurons, blue = DNA. Scale bars = 100um.

**Supplementary Figure 2:**
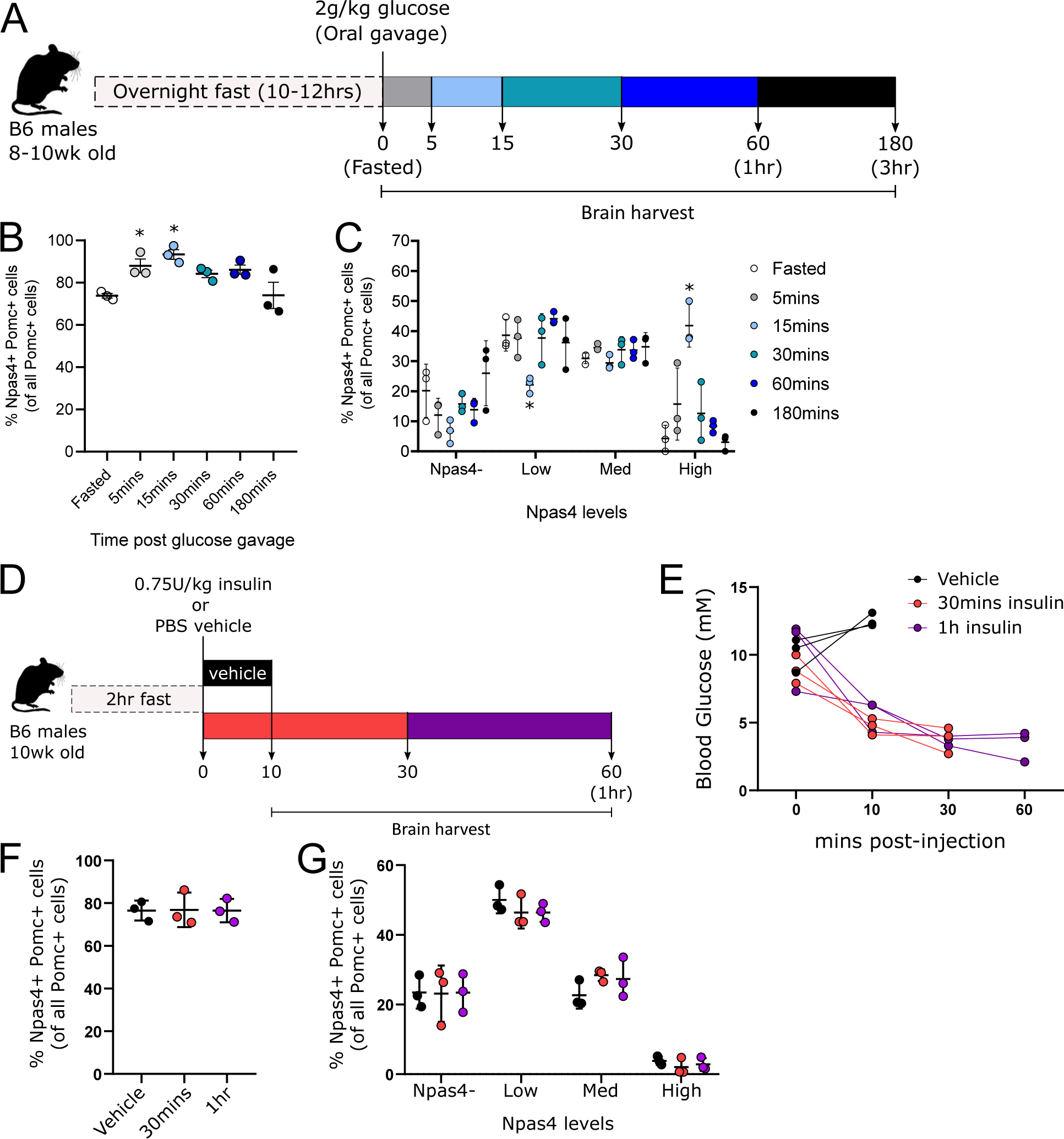
Oral glucose, but not insulin, induces Npas4 in POMC neurons. A) Schematic of timed brain harvests following an oral glucose bolus after a fast. B) Quantifications of Npas4+ Pomc+ cells as a percentage of all Pomc+ cells from RNAscope images obtained from mice at each time point. C) Breakdown of quantifications (B) into bins of Npas4 expression: Npas4- = 0 spots/cell, Low = 1-3 spots/cell, Med = 4-9 spots/cell, High = 10 or more spots/cell. Values are expressed as a percentage of Npas4+ Pomc+ cells in all Pomc+ cells. D) Schematic of timed brain harvests following an intraperitoneal insulin injection. E) Blood glucose measurements following insulin injection. F-G) Quantifications of RNAscope images (F) and breakdown (G) in the same manner as B-C. Data shown as mean with SD. One-way ANOVA with Tukey’s multiple comparisons: *P < 0.05 vs Fasted.

**Supplementary Figure 3:**
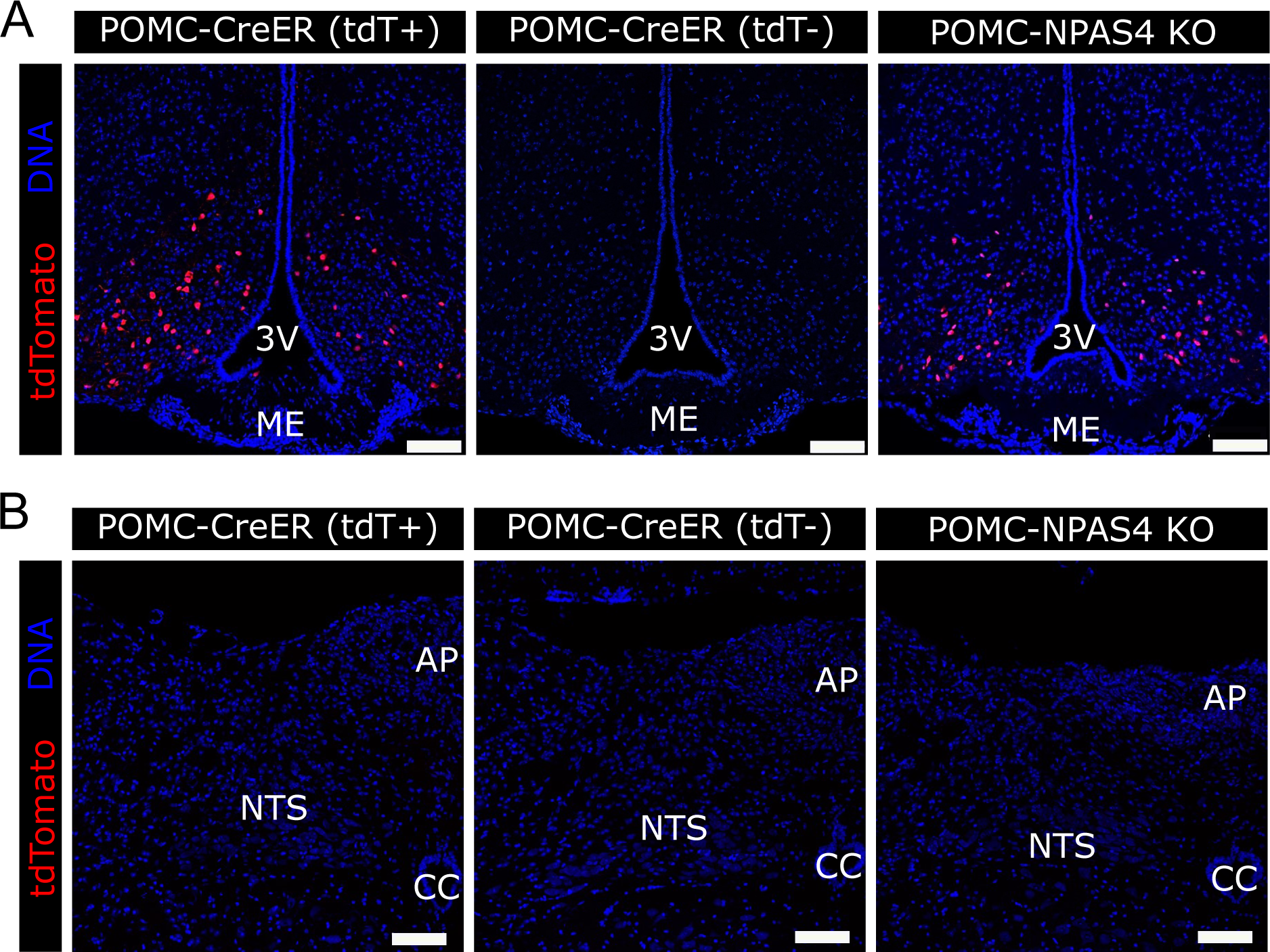
POMC-CreER recombination is restricted to the arcuate nucleus. A-B) Representative sections of the arcuate nucleus (A) and hindbrain NTS regions (B) from tdTomato+ (tdT+) and tdT- POMC-CreER controls and POMC-NPAS4 KO mice, 2 weeks after tamoxifen administration. Red = tdTomato, blue = DNA. 3V = 3rd ventricle, ME = Median Eminence, AP = Area Postrema, NTS = Nucleus of the Solitary Tract, CC = Central Canal. Scale bars = 100um.

**Supplementary Figure 4:**
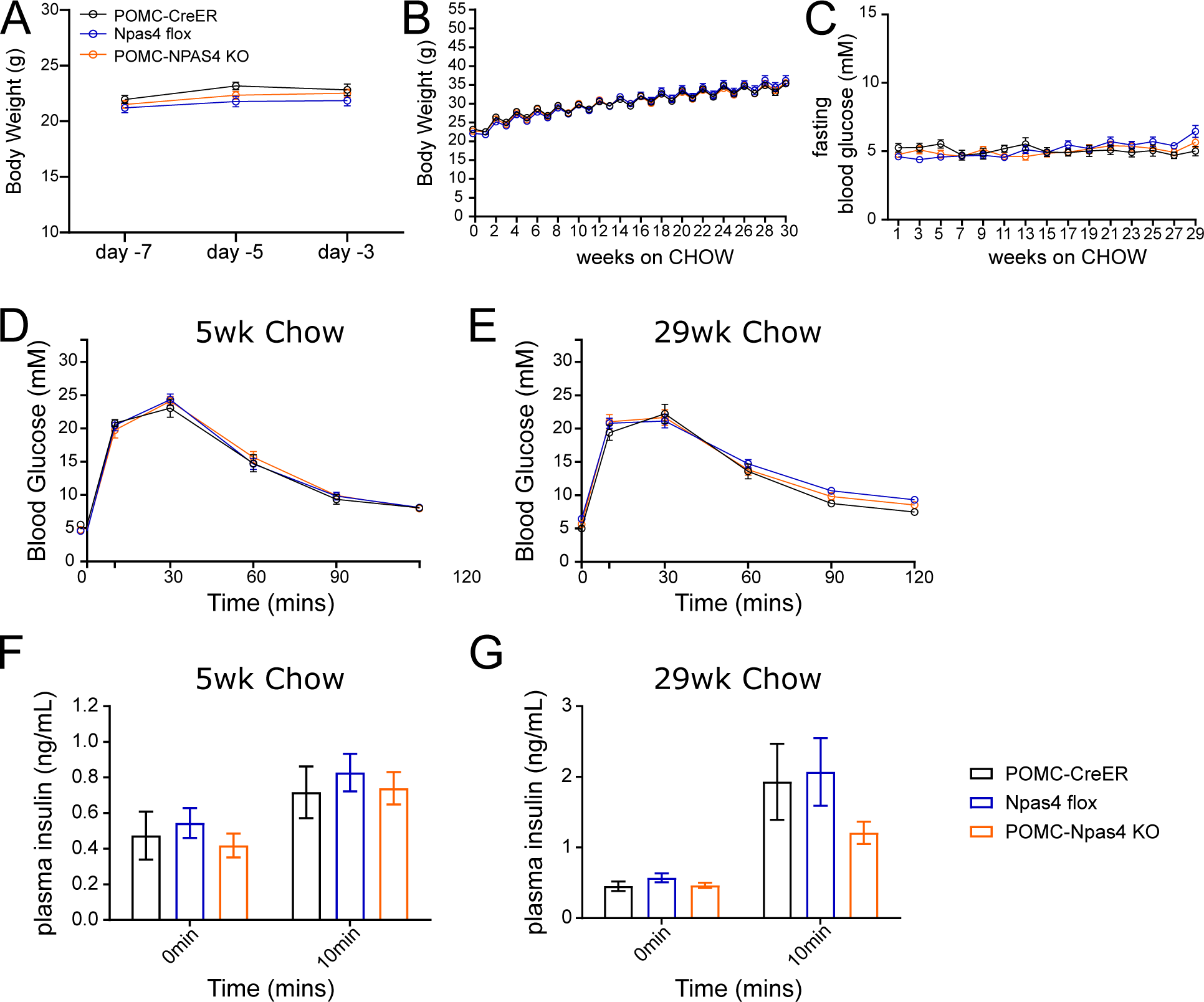
POMC-NPAS4 KO mice show normal body weight gain and glycemia when fed chow. A) Body weights measured before (day-7) and during (days -5, -3) tamoxifen administration to POMC-CreER controls (n=10), Npas4 flox controls (n=22), and POMC-NPAS4 KO mice (n=19). B) Body weights measured weekly during 30 weeks of chow diet after tamoxifen administration. C) Biweekly overnight fasted blood glucose measurements during 30 weeks of chow diet after tamoxifen administration. D-E) Blood glucose measurements during 2g/kg oral glucose tolerance tests performed at 5 weeks (D) and 29 weeks (E) of chow diet. F-G) Plasma insulin measured at 0 and 10 minutes of the oral glucose tolerance tests at 5 weeks (F) and 29 weeks (G) of chow diet.

**Supplementary Figure 5:**
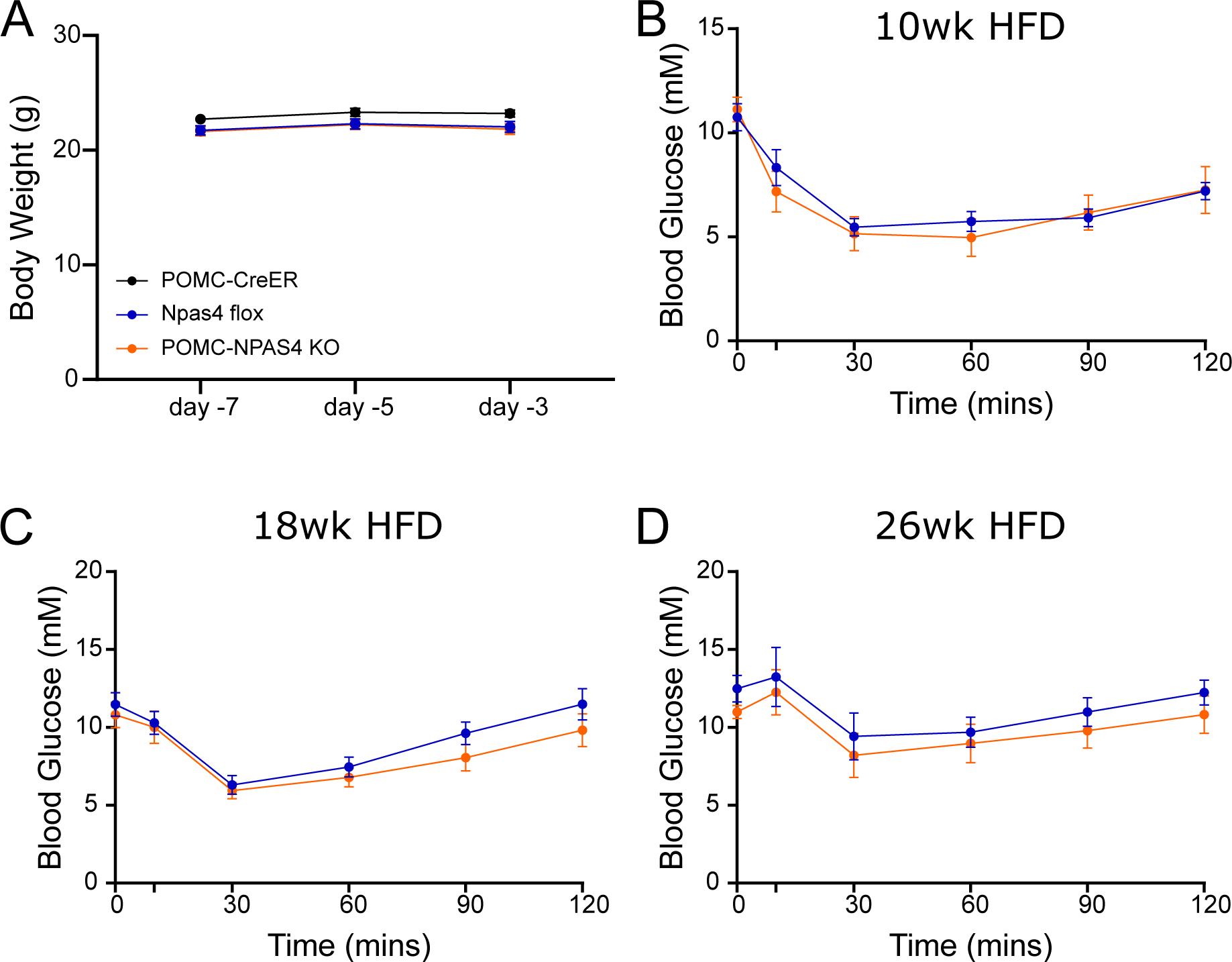
POMC-NPAS4 KO mice show normal glycemia and insulin tolerance on HFD. A) Body weights measured before (day -7) and during (days -5, -3) tamoxifen administration to POMC-CreER controls (n=9), Npas4 flox controls (n=20), and POMC-NPAS4 KO (n=19) mice. B-D) Blood glucose measurements from intraperitoneal insulin tolerance tests performed on Npas4 flox (n=7) and POMC-NPAS4 KO (n=7) mice at 10 weeks (B), 18 weeks (C), and 26 weeks (D) of HFD. 2-way ANOVA with Tukey’s multiple comparisons.

**Supplementary Figure 6:**
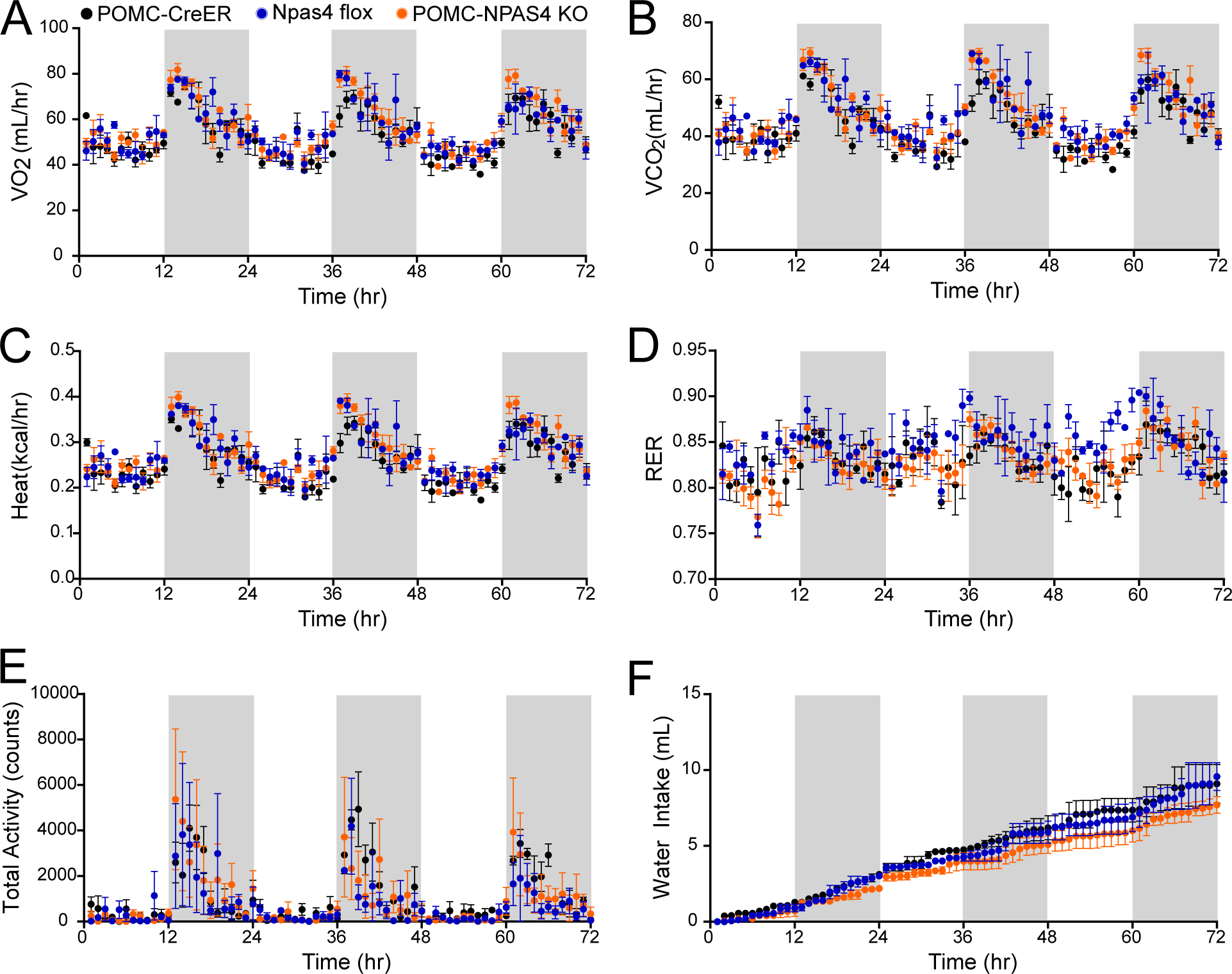
POMC-NPAS4 KO mice show normal indirect calorimetry readouts, activity, and water intake at 6 weeks of HFD. A) Oxygen consumption (VO_2_) measurements (mL/hr) for POMC-CreER (n=3), Npas4 flox (n=2), and POMC-NPAS4 KO (n=3) mice. B) Carbon dioxide production (VCO_2_) measurements (mL/hr) for all genotypes. C) Energy expenditure as heat production estimated from VO_2_ (kCal/hr) for all genotypes. D) Respiratory exchange ratio (RER) for all genotypes. E) Locomotor activity over time in counts for all genotypes. F) Water intake measurements (mL) over time in all genotypes.

**Supplementary Figure 7:**
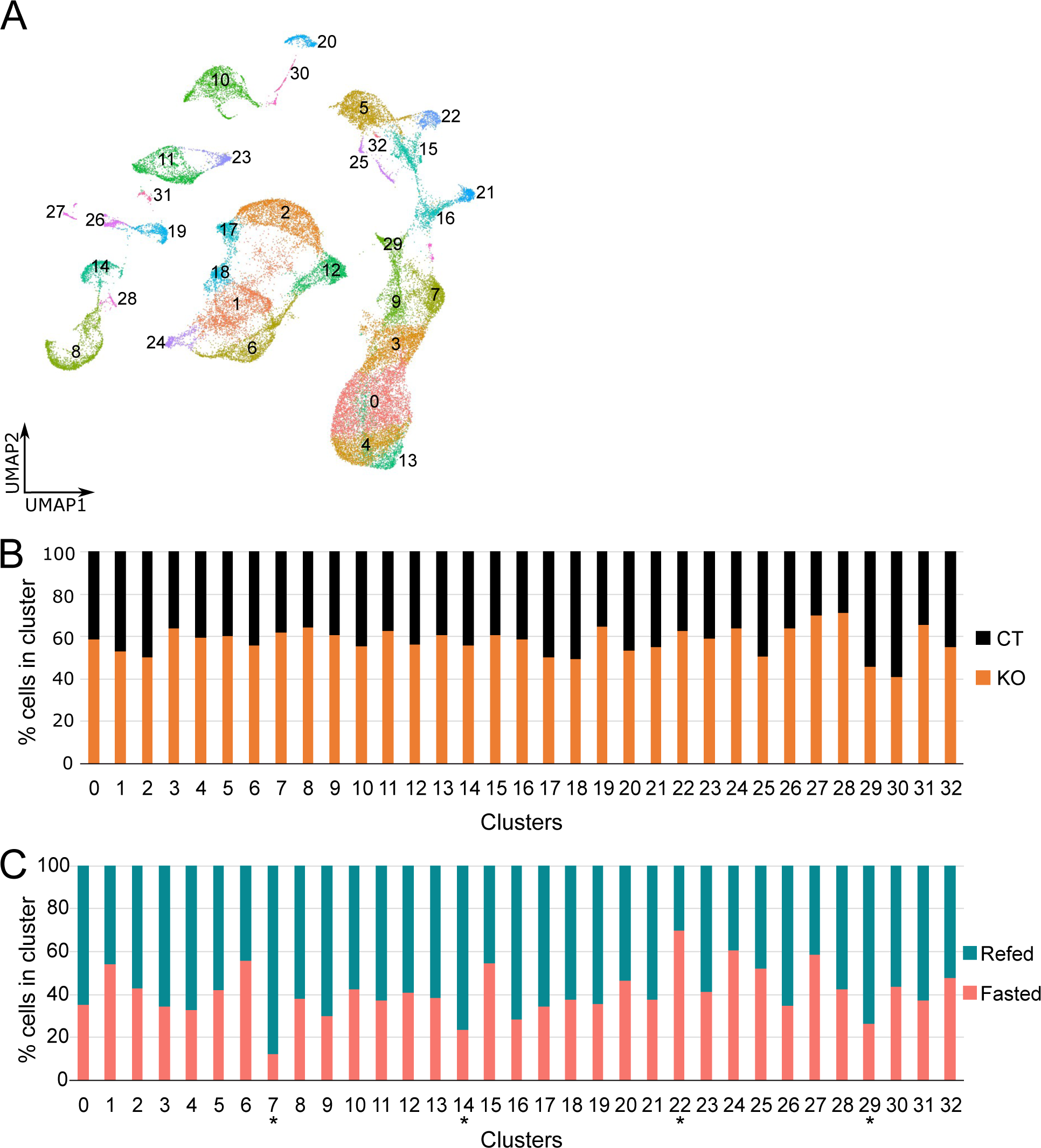
Clustering of mouse arcuate nucleus scRNA-seq data is not driven by genotypes. A) All samples integrated and projected as 32 clusters in dimensionally reduced UMAP space. Clusters are labelled with their original Seurat-assigned cluster numbers, in order of cluster size, with cluster 0 being the largest. B-C) Proportion bar graphs showing the percentage of cells in each cluster from CT and KO samples (B) or Refed and Fasted samples (C). Asterisks on clusters 7, 14, 22, and 29 mark these clusters as having a skewed proportion in favour of cells from Refed or Fasted samples.

**Supplementary Figure 8:**
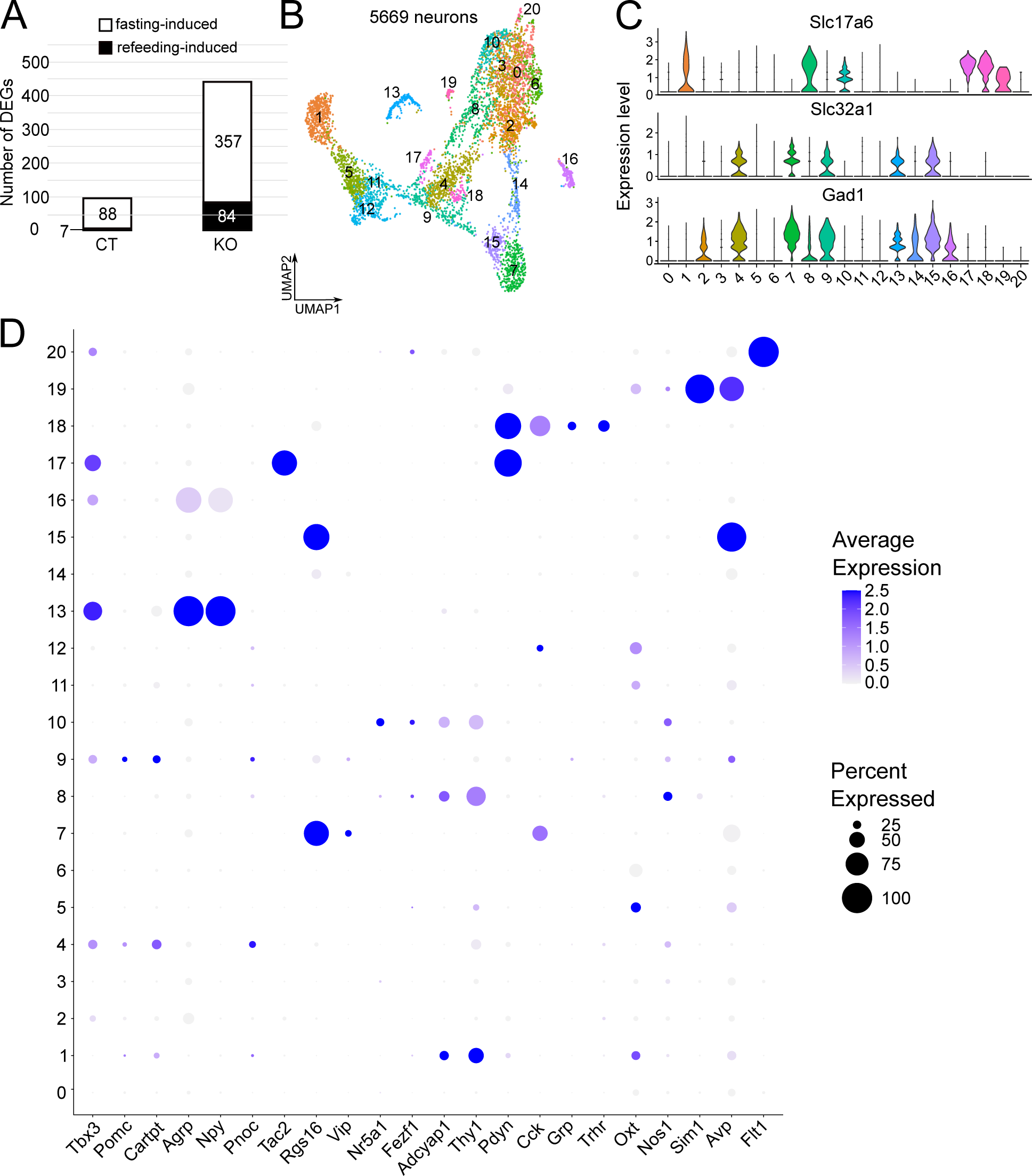
Identification of reclustered neurons. A) Number of DEGs between fasting and refed POMC neurons in CT and KO. B) Reclustered 5669 neurons projected in UMAP space as 21 clusters. C) Violin plots showing expression of glutamatergic neuron marker gene *Slc17a6* and GABAergic neuron marker genes *Slc32a1* and *Gad1*. D) Dotplot showing average expression of known marker genes for neuronal subpopulations in various hypothalamic nuclei. Dot sizes represent percent of cells in each cluster that express a certain gene on the x-axis (“Percent Expressed”). Increasing shades of blue represent average expression levels in the cluster (“Average Expression”), with darker shades of blue being higher average expression.

**Supplementary Figure 9:**
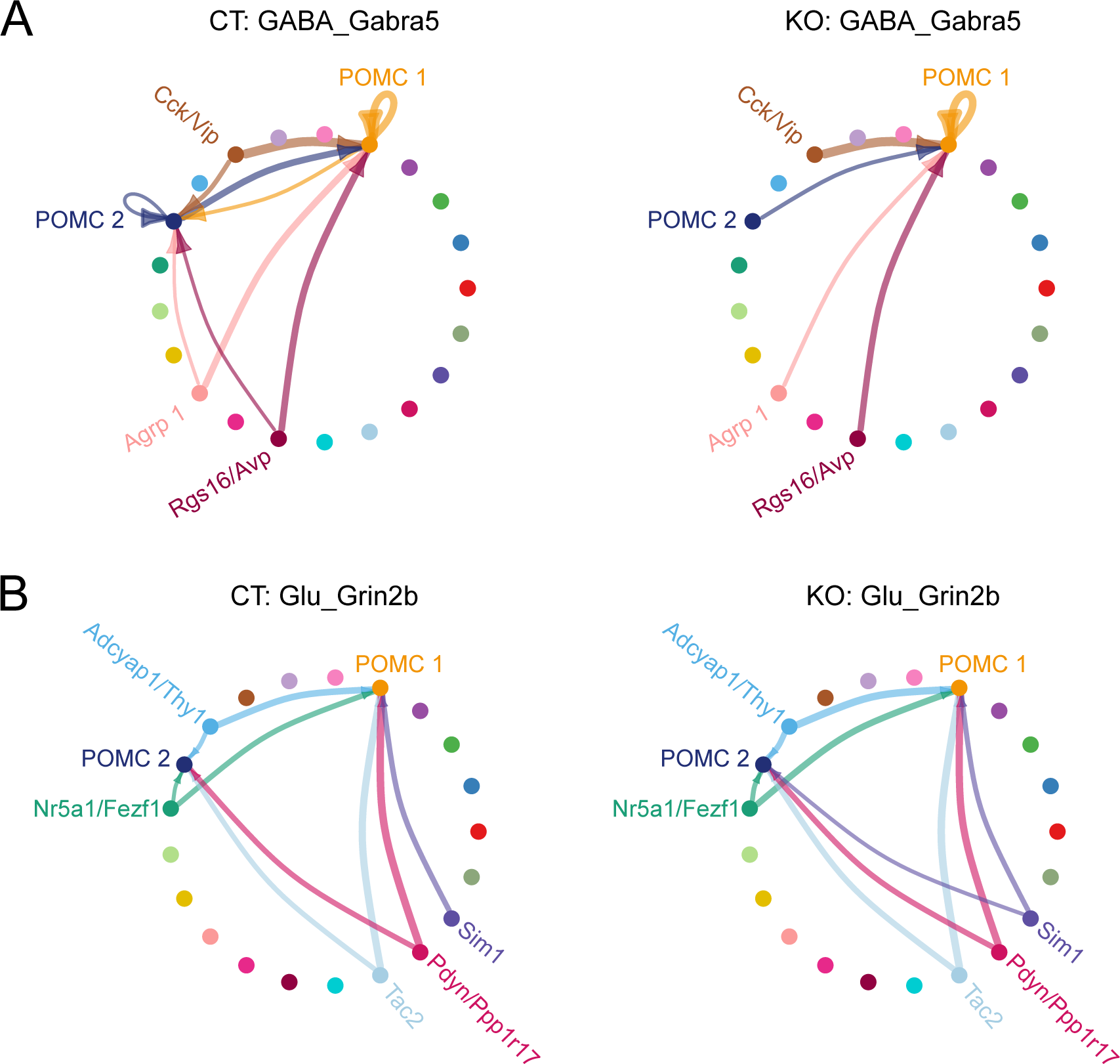
POMC neurons with NPAS4 KO lose GABAergic and gain glutamatergic inputs from other regions. Circle plots showing individual signal-target interactions between all sender regions to POMC neurons only. Each cluster is represented as a coloured dot on the periphery of the circle, with arrows indicating the direction of communication and the significance of the interaction as the thickness of each arrow. Plots are shown for A) GABA_Gabra1 and B) Glu_Grin2b between POMC-CreER and POMC-NPAS4 KO neuron clusters.

## Notes

### Competing Interest Statement

The authors have declared no competing interest.

