## Supplemental methods for "NPAS4 is an allostatic regulator of POMC neuronal activity during diet-induced obesity"

### Supplementary Methods

#### Brain harvests and preparation for histology

Mice were anesthetized with isoflurane under presence of $O_2$ and euthanized by asphyxiation with $CO_2$. After visually confirming that breathing had stopped, the mouse was removed from the isoflurane chamber, sprayed with 70% ethanol, and dissected to expose the liver and stomach in the abdominal cavity within one minute of removal from the chamber. The carcass was dissected to expose the heart, and the right atrium was punctured. A 27½ gauge needle with tubing (1mm diameter) attached to a syringe was inserted into the left ventricle, and the carcass was flushed of blood with 5mL of cold PBS. The 5mL syringe was replaced with a 30mL syringe containing 20mL of 4% paraformaldehyde (PFA), and the carcass was transcardially perfused. The perfused carcass was decapitated, the head was flayed to expose the skull, and the skull was removed until the intact whole brain could be removed.

The next morning, the PFA was removed, and brains were washed three times with PBS, 5 minutes each, and transferred to a clean tube containing 10mL of 20% sucrose (w/v solution in water) to submerge the tissue for 24 hours, or until the brain had sunk to the bottom of the tube, at 4°C. The 20% sucrose solution was removed and replaced with 10mL of 30% sucrose (w/v solution in water) to submerge the tissue for 24 hours, or until the brain had sunk to the bottom of the tube, at 4°C. Next, the brain was removed from the sucrose solution, transferred to a small dish containing OCT cryomatrix (Epredia; ref 6502), and thoroughly coated in OCT using blunt-ended forceps. The brain was bisected coronally at the level of the caudal edge of the hypothalamus, visible on its ventral surface, and the caudal portions of tissue were discarded. The remaining brain tissue was embedded in a cryomold with fresh OCT and frozen within the cryomold on a bed of dry ice. The frozen brains were stored at -80°C until sectioning. Prior to cryosectioning, embedded brains were equilibrated to -20°C for at least 2 hours. The cryostat was set to -20°C and the embedded brains were trimmed at 50µm until tissue became visible. Sections were collected directly onto Superfrost Plus glass slides and stored at -80°C. Once sectioning was complete, blocks were re-sealed and re-frozen with OCT cryomatrix before storing at -80°C.

**RNAscope fluorescent *in situ* hybridization**

Briefly, cryosections were washed in PBS to remove OCT, baked in the HybEZ oven (ACDbio) at 60°C for 1 hour to improve tissue adherence, and placed into target retrieval buffer (ACDbio; 322000) heated to 95-98°C in a steamer for five minutes before being immersed in 100% ethanol for three minutes. A hydrophobic barrier was drawn around the sections with the ImmaEdge pen (Vector Laboratories; H-4000), and sections were incubated with protease III (ACDbio; 322337) in the HybEZ oven at 40°C for 30 minutes. Sections were washed with $dH_2O$ and incubated with probes in the HybEZ oven at 40°C for 2 hours. Sections were washed with 1X wash buffer (ACDbio; 310091) and left overnight in 5X SSC buffer, diluted in water from 20X SSC buffer (Ambion; AM9763), at room temperature. The next day, sections were washed with wash buffer and underwent amplification and development of each channel signal with channel-specific HRP reagents and Opal dyes. After a 5 minute incubation with provided DAPI, slides were washed and mounted with Diamond antifade mounting media and stored at 4°C until imaged with a Leica SP8 confocal microscope.

**RNAscope quantification**

For all experiments, three sections of the ARC were quantified per mouse. For each RNAscope experiment, the technical positive control section was used to confirm successful protocol completion, and the technical negative control section was used to determine the imaging settings to prevent any background signal in any channel. The left and right sides of the arcuate nucleus were imaged as Z-stacks separately for each section using the Leica SP8 confocal microscope 20X objective. The resulting image files were converted to .ims file formats and quantified with Imaris v9 (Oxford Instruments). The Cells with Vesicles feature was used on the image, specifying Cells as the areas with *Pomc* signal that marked the neuronal cell bodies, and specifying Vesicles as the *Npas4* punctate signals. For each image, the threshold for Cells was manually set to completely cover all areas with detectable *Pomc* signal, and cell boundaries were automatically calculated by the software when cell diameter was set to 10μm. Cells that had detectable *Pomc* signal but no nucleus were manually removed from the final quantifications. Vesicle diameter was set to 2μm, and thresholds for Vesicles was manually set to count all *Npas4* punctate signals within the regions covered by *Pomc* signal. Final statistics reports were exported as counts of vesicles per cell, and exported data was manually organized on Microsoft Excel to quantify the number of *Npas4* spots per *Pomc*+ cell and the proportion of *Npas4+/Pomc+* cells. Due to the relatively low numbers of *Npas4* puncta in the ARC, *Npas4+* cells were classified into 4 levels of *Npas4* expression based on the number of spots in a cell of interest: *Npas4*- (0 spots), *Npas4*-low (1-3 spots/cell), *Npas4*-medium (4-9 spots/cell), or *Npas4*-high (10 or more spots/cell).

**Immunostaining**

Fixed frozen sections of mouse brain were washed for 5 minutes in PBS to remove OCT cryomatrix before permeabilizing for 7 minutes with 1% SDS. Sections were washed three times with PBS, 5 minutes each. A hydrophobic barrier was drawn around each tissue section with a pap-pen (Cedarlane; MU22-A) and the sections were blocked with 5% horse serum in PBS for 30 minutes at room temperature in a humidified staining box. After blocking, the sections were washed three times with PBS, 5 minutes per wash. Primary antibodies were diluted in PBS with 5% horse serum and sections were incubated with primary antibodies overnight at 4°C. Primary antibodies used were anti-NPAS4 (Activity Signalling AS-AB18A-100; 1:500) and anti-$GABA_AR$ β3 (Synaptic Systems 224 403; 1:500).

The next morning, the sections were washed three times with PBS, 5 minutes per wash. Secondary antibody donkey anti-rabbit FITC (Jackson ImmunoResearch 706-166-148; 1:450) and nuclear stain TO-PRO-3 were diluted in PBS with 5% horse serum, and sections were incubated with the dilutions for 1 hour in the dark at room temperature. Sections were washed three times with PBS, 5 minutes per wash, and mounted with Diamond antifade mounting media and glass coverslips. Mounted slides were stored in a slide box at 4°C until they were imaged using a Leica SP8 confocal microscope.

**Mouse hypothalamus dissociation for scRNA-seq**

After an overnight fast with or without 1 hour of refeeding in the morning, mice were anesthetized with isoflurane and euthanized by cervical dislocation. The head and neck were sprayed down with 70% ethanol, and heavy surgical scissors were used to decapitate the mouse. The brain was rapidly removed from the skull, placed onto a petri dish with its ventral side facing upwards, rinsed in 15mL of PBS, and wetted with 5mL of Hibernate® AB complete medium (BrainBits; HAB). A clean razor blade was used to bisect the brain coronally at the caudal edge of the hypothalamus, visible on its ventral surface, and the caudal portions of tissue were discarded. The remaining brain tissue was placed into 7mL of HAB medium and placed on ice until all samples had been collected. Using a clean razor blade, two 1mm-thick coronal sections were collected from each brain and moved to a petri dish with HAB medium. Under a dissection microscope (Olympus; SZX16), the ARC was roughly dissected out from each thick section.

The tissue was placed into a 15mL polystyrene tube (VWR; 21008-212) containing 2mL of sterile-filtered 2mg/mL papain solution, made by dissolving solid papain (BrainBits; PAP) in Hibernate® A medium without calcium (BrainBits; HACA) for 20-30 minutes at 37°C. The tube containing papain and tissue was placed in a shaking water bath (VWR) set to 32°C, and the tissue was dissociated by shaking at 170-180rpm for 40 minutes. The tissue was removed from the papain using a wide-bore pipette tip, transferred to a clean tube containing 2mL of HAB medium, and incubated for 5 minutes on ice. Fire-polished siliconized 9-inch glass Pasteur pipettes were prepared by polishing the tip to 0.6-0.8mm diameter over a bunsen burner and siliconizing with Sigmacote® (Sigma; SL2-100ML). Using the fire-polished pipettes, the tissue was triturated ten times over one minute, then allowed to settle for one minute on ice. The supernatant was transferred to an empty 15mL tube, and the remaining tissue was resuspended in a fresh 2mL of HAB medium. The trituration was repeated 2 more times, for a total of 6mL of dissociated cells. The 6mL of dissociated cell solution was carefully applied to the top of the prepared OptiPrep™ (Sigma; D1556-250ML) density gradient [41], and the gradient was centrifuged at 800g for 15 minutes at 4°C. The top 7mL of the centrifuged gradient, containing debris and oligodendrocytes, were aspirated. The next 2.5mL, excluding the microglia pellet at the bottom, were collected and diluted with 5mL of HAB medium. The cell solution was centrifuged for 2 minutes at 200g at 4°C, and the supernatant was removed. The cell pellet was resuspended in 1mL of PBS- (cytiva; SH30028.02) and filtered through a 30µm strainer (Miltenyi Biotec; 130-041-407) into a clean tube. An additional 500µL of PBS- was used to rinse the tube and this was also filtered. The cell solution was centrifuged for 2 minutes at 200g at 4°C, the supernatant was removed, the cells were resuspended in 100µL of PBS-, and transferred to a nuclease-free 1.5mL microcentrifuge tube, which was kept on ice. 10µL of the single cell solution was stained with 10µL of trypan blue dye, and 10µL of this mixture was visualized on a hemacytometer to confirm successful dissociation with minimal debris and high cell viability. A final cell count was calculated by manually counting the total number of bright visible cells in all quadrants, calculating the average number of cells per quadrant, and multiplying by $10^4$, the dilution factor, and the resuspension volume in mL.

**Sequencing**

Two to four samples' libraries were diluted to 4nM and pooled before sequencing, depending on chronological order of mouse availability. All libraries were sequenced with the Illumina NextSeq500 on manual mode using a 75-cycle High Output v2.5 kit (Illumina FC-404-2005), using the following cycle numbers: 8 for index 1 (i7), 0 for index 2 (i5), 28 for read 1, and 56 for read 2. For libraries generated using Chromium Next GEM v3.1 kits, cycle numbers were: 10 for index 1 (i7), 10 for index 2 (i5), 28 for read 1, and 44 for read 2. Sequencing was repeated for all libraries until all samples reached a minimum read depth of 20000 reads/cell as determined by downstream processing of sequencing data.

**Analysis of scRNA-seq data**

Binary base call (BCL) files of sequencing data were obtained as outputs from the Illumina NextSeq500. Cell Ranger v7.0 (10x Genomics) was used for processing of all scRNA-seq data. FASTQ files from sequencing outputs with *cellranger mkfastq*. Alignment, demultiplexing, filtering, barcode counting, and UMI counting was performed with *cellranger count* separately for each sample, combining multiple sequencing runs' worth of FASTQ files for the same sample. FASTQ files were aligned to the reference mouse genome GRCm38. Reads/cell, genes/cell, and number of cells per sample were obtained at the end of the Cell Ranger pipeline.

Outputs of Cell Ranger were carried forward to the R package Seurat [42] v4.0.4. First, Seurat objects were created from the Cell Ranger outputs and cells that expressed more than 20% mitochondrial genes or more than 6000 genes/cell were filtered out. The metadata was edited to include mouse number (sample ID), fasted or refed status, and genotype. Next, the objects were normalized with SCTransform and integrated to form one Seurat object with all samples, and clustered in UMAP space at a resolution of 0.6. The biological identities of clusters were manually determined with known marker gene expression. Differentially expressed gene (DEG) analysis was performed using *FindMarkers* or *FindAllMarkers* functions when needed. For neuronal cells in the dataset that were subsetted from the original Seurat object containing all cell types, additional clustering was performed at a resolution of 1.5 to address the high degree of heterogeneity. Following identification of cluster markers and identities, cell-cell communication analysis was performed with NeuronChat [46]. Data matrices from the SCT assay's "data" slot and metadata were extracted for all neurons in order to preserve labelling at the cluster level.
